# *Amaranthus viridis*-derived phytopriming reprograms redox homeostasis and limits arsenic accumulation in rice

**DOI:** 10.64898/2026.08.06.743420

**Authors:** Susmita Poddar, Sanket Roy, Anwesha Behera, Ishanee Das Sharma, Surupa Chakraborty, Rajib Sengupta, Natasha Das, Surajit Bhattacharya

## Abstract

Arsenic (As) poses a major threat to rice productivity and food safety due to its high bioaccumulation potential and subsequent entry into the human food chain. In rice, As impairs seed germination, disrupts morpho-anatomical development, and induces oxidative stress. This study evaluates seed priming with an aqueous extract of the agricultural weed *Amaranthus viridis* (AvE) as a sustainable strategy to alleviate As-induced phytotoxicity. AvE priming significantly improved germination (71–75%) and morpho-physiological performance under As stress. It reduced oxidative stress markers, including H_2_O_2_ (21–38%), malondialdehyde (13–26%), and proline (18.9–44.7%), while increasing antioxidant metabolites, polyphenols and glutathione by up to 2.34-fold and 41%, respectively. Microscopy confirmed restoration of cellular integrity and anatomical organisation in primed seedlings. ICP-OES analysis showed that AvE priming reduced root As uptake by up to 39%, root-to-shoot translocation by up to 58%, and grain As accumulation by up to 95% compared with unprimed plants. qRT-PCR revealed modulation of genes involved in As homeostasis, indicating coordinated physiological and transcriptional responses. Importantly, improved agronomic performance further demonstrated the translational potential of this approach. This study provides the first evidence that *A. viridis* extract is a cost-effective, sustainable biostimulant for producing low-As rice in contaminated regions.

## 1. Introduction

Rice (*Oryza sativa* L.) serves as the principal staple food for a large proportion of the global population, and it remains indispensable in countries such as India, where it underpins food security and livelihoods. In India, with a population exceeding 1.4 billion, nearly two-thirds of the population depends on rice as a primary dietary component due to its wide availability, affordability, and nutritional value, particularly as a rich source of carbohydrates (Birla et al. 2017). Consequently, rice is often regarded as the backbone of sustenance for both rural and urban populations (Mahajan et al. 2017). To meet this demand, India has emerged as the second-largest rice producer globally, with over 43 million hectares under cultivation (Varma 2017). However, the sustainability of rice production is increasingly threatened by arsenic (As) contamination, particularly in regions where groundwater used for irrigation contains elevated levels of this toxic metalloid. The use of As-contaminated groundwater in paddy cultivation has led to widespread accumulation of arsenic in agricultural soils, making irrigation practices a primary route of As entry into rice plants (Rahman et al. 2014; Awasthi et al. 2017). Notably, rice exhibits a significantly greater propensity for arsenic accumulation compared to other cereal crops, primarily due to its cultivation under flooded conditions and the activity of endogenous transport systems that facilitate arsenic uptake (Williams et al. 2007). This accumulation adversely affects plant growth, germination, and physiological processes, while also posing severe health risks to humans through dietary exposure (Murugaiyan et al. 2021). Alarmingly, arsenic contamination in rice has emerged as a major concern across the Asiatic Plain, particularly in countries such as Bangladesh, China, Nepal, and India (Ayers et al. 2019), placing an estimated 230 million people at risk of chronic arsenic poisoning, including approximately 180 million in Asia (Shaji et al. 2021).

Given the highly toxic and carcinogenic nature of arsenic, mitigating its accumulation in rice grains has become a critical priority for ensuring food and nutritional security. Several approaches, including genetic and biotechnological interventions targeting key transporters and detoxification pathways, have been explored (Zhao et al. 2010). However, none of these strategies has successfully achieved the production of arsenic-free rice grains. A major limitation lies in the inherent lack of selectivity of plant transport systems, which inadvertently facilitate arsenic uptake alongside essential nutrients (Das et al. 2020). Moreover, the widespread deployment of genetically modified (GM) rice remains constrained due to regulatory stringency and limited societal acceptance in many parts of the world. These challenges highlight the urgent need for alternative, non-transgenic, and sustainable approaches to reduce arsenic accumulation in rice.

Seed priming has emerged as an effective, low-cost, and environmentally sustainable strategy to enhance plant tolerance against various abiotic stresses. This pre-sowing treatment involves controlled hydration of seeds, which activates metabolic processes associated with germination, leading to improved seedling establishment and vigor. Importantly, priming has been shown to enhance antioxidant defences, promote DNA repair, and regulate stress-responsive pathways, thereby mitigating oxidative damage induced by abiotic stressors such as arsenic (Basra et al. 2005). Despite its proven effectiveness in addressing multiple stress conditions, the application of seed priming for mitigating arsenic toxicity and accumulation in rice remains relatively underexplored.

In parallel, plant-derived phytochemicals have attracted significant attention due to their potent antioxidant properties and their role in mitigating oxidative stress. Weeds, often considered undesirable in agricultural systems, are in fact rich sources of diverse bioactive compounds that confer resilience under harsh environmental conditions. These phytochemicals, including polyphenols, flavonoids, saponins, and terpenoids, play essential roles in scavenging reactive oxygen species (ROS) and maintaining cellular homeostasis. Beyond their agricultural relevance, such compounds have also been widely studied for their therapeutic potential, including roles in managing neurodegenerative disorders, cardiovascular diseases, and microbial infections (Ghosh et al. 2015). However, their application in crop stress management, particularly through seed priming approaches, remains limited.

*Amaranthus viridis* L., a widely distributed weed belonging to the family Amaranthaceae, is known for its rich phytochemical composition and diverse medicinal properties. Commonly found across tropical and subtropical regions, including Asia, North America, and Europe, this species has traditionally been used for its antipyretic, analgesic, and anti-inflammatory properties (Reyad-ul-Ferdous et al. 2015; Eluwa, 1977). Phytochemical analyses have revealed the presence of key bioactive constituents such as alkaloids, phenolics, flavonoids, saponins, tannins, steroids, and triterpenoids, along with essential amino acids. Owing to this diverse biochemical profile, *A. viridis* exhibits strong antioxidant, antimicrobial, and therapeutic potential (Reyad-ul-Ferdous et al. 2015). Given its rich reservoir of antioxidant phytochemicals, it is hypothesized that aqueous extracts of *A. viridis* can be effectively utilized as a priming agent to mitigate arsenic-induced oxidative stress in rice. Such a phytopriming approach may enhance stress tolerance by modulating ROS scavenging systems and activating defence-related signalling pathways. Therefore, the present study aims to evaluate the potential of *A. viridis*-based seed priming (AvE) in alleviating arsenic toxicity and reducing its accumulation in rice seedlings. By integrating physiological, biochemical, anatomical, and molecular analyses, this work seeks to establish a novel, eco-friendly, and non-transgenic strategy for improving arsenic stress tolerance and ensuring safer rice production in contaminated agroecosystems.

## 2. Materials and Methods

### 2.1 Collection of *Amaranthus viridis* and preparation of aqueous extract

Fresh plants of *Amaranthus viridis* were collected from the vicinity of Amity University Kolkata, Newtown, West Bengal, India. Shoot tissues were separated, thoroughly washed under running tap water to remove adhered debris, and blotted dry. The samples were subsequently air-dried at 37 °C for 10–12 days in a hot-air incubator until constant weight was achieved. Dried tissues were ground into a fine powder using a sterile mortar and pestle. Aqueous extraction was performed by suspending the powdered biomass in distilled water at a concentration of 10 mg mL□¹, followed by continuous stirring. The extract was filtered to remove insoluble debris and sterilised using a 0.2 µm syringe filter. The resulting filtrate was used for seed priming and phytochemical screening.

### 2.2 Seed priming treatment and experimental design

Seeds of rice (*Oryza sativa* L. cv. IR64) were surface sterilized following the protocol of Das et al. (2017). The sterilized seeds were subjected to three treatment conditions: (i) unprimed, (ii) priming with 5 mg mL□¹ *A. viridis* extract (AvE), and (iii) priming with 10 mg mL□¹ AvE. For priming, 100 seeds were incubated in 10 mL of the respective solutions for 24 h in the dark at 28 °C with gentle agitation (80 rpm). Following incubation, seeds were rinsed thoroughly with sterile distilled water and air-dried on sterile blotting paper prior to sowing. To assess arsenic stress, sodium arsenate [As(V)] was incorporated into water agar medium at a final concentration of 10 mg L□¹. Four experimental groups were established: (i) Control (unprimed seeds without As), (ii) As(V) + unprimed, (iii) As(V) + AvE 5 mg mL□¹, and (iv) As(V) + AvE 10 mg mL□¹. Fifteen seeds were sown per Petri plate, and each treatment was performed in triplicate. Plates were incubated in darkness at 28 °C for 3 days, followed by growth under a 16 h light/8 h dark photoperiod until 14 days after sowing (DAS). Seedlings were then either harvested for further analyses or grown to maturity under continuous exposure to As(V) stress.

### 2.3 Qualitative phytochemical screening

Qualitative phytochemical analyses of the AvE extract were conducted using standard protocols to detect major classes of bioactive compounds, including flavonoids, phenolics, tannins, saponins, terpenoids, proteins, volatile oils, anthocyanins, and betacyanins (Harborne. 1984; Evans. 2009; Brain and Turner. 1975; Vinoth and Manivasagaperumal, 2012) (Supplementary Material 1). The phytochemical profiling of the samples was performed using an Agilent 6530 Liquid Chromatography–Quadrupole Time-of-Flight Mass Spectrometer (LC/Q-TOF-MS, Agilent Technologies, Santa Clara, CA, USA). Mass spectrometric detection was carried out using an electrospray ionization (ESI) source operated in positive and/or negative ionization mode. The Agilent 6530 Q-TOF analyzer provides high-resolution accurate-mass measurements with mass accuracy below 2 ppm and a mass range of m/z 20–20,000. The obtained mass spectra were processed for tentative identification of metabolites and other bioactive compounds present in the sample (Arthur et al. 2017).

### 2.4 Germination parameters and seedling growth analysis

Germination was monitored at 3, 7, and 14 days after sowing (DAS), and seeds with radicle emergence ≥2 mm were considered germinated. Germination percentage was calculated as:

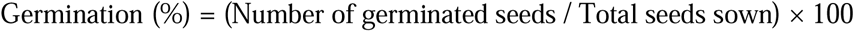

At 14 DAS, root length, shoot length, fresh weight (FW), and dry weight (DW) were recorded. Dry weight was determined after oven drying at 80 °C until constant weight.

Seedling vigor indices were calculated following Abdul-Baki and Anderson (1973):

- Vigor Index I (VI-I) = Germination (%) × Seedling length (root + shoot)
- Vigor Index II (VI-II) = Germination (%) × Seedling dry weight

### 2.5 Relative water content (RWC)

Relative water content was determined following Barrs and Weatherley (1962). The FW of seedlings was recorded, followed by incubation in distilled water for 24 h to obtain turgid weight (TW). Samples were then oven-dried at 80 °C for 24 h to obtain DW. RWC was calculated as:

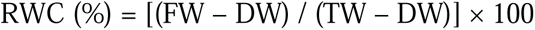

### 2.6 Chlorophyll estimation

Chlorophyll content was estimated using dimethyl sulfoxide (DMSO) extraction (Barnes et al. 1992). Approximately 20 mg of shoot tissue was incubated in 5 mL DMSO at 65 °C until complete pigment extraction. Absorbance was recorded at 664.9 nm and 648.2 nm, and chlorophyll concentrations were calculated using standard equations.

### 2.7 Biochemical assays for oxidative stress and antioxidants

Hydrogen peroxide (H□O□) content was measured following Velikova et al. (2000), while lipid peroxidation was assessed via the Thiobarbituric Acid Reactive Substances (TBARS) assay (Heath and Packer, 1968). Proline content was quantified using the method of Bates et al. (1973). Total phenolic content was determined using the Folin–Ciocalteu method with gallic acid as a standard. To estimate total flavonoid content, the aluminium chloride colorimetric method is used and the total flavonoid content was expressed in mg Quercetin equivalent/g fresh weight (Zhishen et al, 1999). Reduced glutathione (GSH) levels were measured following Anderson (1985).

### 2.8 Light microscopy for anatomical analysis

Transverse sections (TS) of root tissues (1 cm away from the root–shoot junction) were prepared from 14 DAS seedlings. Sections were stained with 0.25% (w/v) safranin for 1 min, mounted in 10% glycerol, and observed under a light microscope to examine anatomical features.

### 2.9 Scanning electron microscopy (SEM)

For ultrastructural analysis, root sections were fixed in 2.5% glutaraldehyde, followed by dehydration through a graded ethanol series. Samples were sputter-coated with gold– palladium and visualised using a scanning electron microscope Carl Zeiss EVO LS10 following standard protocol (Ray et al. 2022). Elemental analysis and mapping of the root transverse sections were performed using an energy-dispersive X-ray analysis (EDAX) detector coupled to the SEM to determine the presence of As and its distribution in the rice vascular bundle.

### 2.10 Determination of arsenic content (ICP–OES)

Total arsenic concentration in root and shoot tissues of 14 DAS seedling along with grain of mature rice plants were quantified using inductively coupled plasma optical emission spectrometry (ICP–OES; PerkinElmer). Dried samples (100 mg) were digested with concentrated nitric acid, followed by hydrogen peroxide treatment. The digested samples were filtered (0.2 µm) prior to analysis.

The translocation factor (TF) was calculated as: TF = (As concentration in shoot / As concentration in root) × 100

### 2.11 Quantitative real-time PCR (qRT-PCR)

Total RNA was extracted from root tissues using the PureLink RNA Mini Kit (Thermo Scientific). First-strand cDNA was synthesized using SuperScript™ First-Strand Synthesis System. Gene expression analysis was performed using qRT-PCR, with Histone H3 as the internal control. Target genes included *OsABCC1*, *OsARM1*, *OsHAC1;1*, *OsLSI1*, *OsNRAMP1*, *OsPT1*, *OsPT8*, *OsCLT1*, *OsRBOHC*, and *OsSHR2*. Primer details are provided in Supplementary Material 5.

### 2.12 Agronomic evaluation under field conditions

Field experiments were conducted to evaluate the agronomic performance of primed and unprimed plants grown to maturity under standard cultivation practices. Parameters recorded included days to flowering, plant height, number of productive tillers, panicle length, number of grains per panicle, and grain weight. Data were collected from five randomly selected plants per replication (Bheemanahalli et al. 2017, Huang et al. 2020, Singh et al. 2024).

### 2.13 Statistical analysis

All experiments were conducted with three biological replicates (n = 3). Data were expressed as mean ± standard deviation (SD). Statistical significance was determined using one-way ANOVA followed by Tukey’s HSD test at *p* ≤ 0.05 using GraphPad Prism.

## 3. Results

### 3.1 Metabolite profiling of *A. viridis* extract (AvE) by LC–MS

Qualitative LC–MS analysis of the aqueous extract of *Amaranthus viridis* (AvE) revealed a chemically diverse metabolite profile comprising more than 37 compounds (Table 1). The identified metabolites were categorised into multiple functional classes, including antioxidants, phytohormones, osmoprotectants, lipid derivatives, organic acids, and stress-associated secondary metabolites. Notably, several potent antioxidant compounds were detected, including ascorbic acid, α-tocopherol, caffeic acid derivatives, flavonoid glycosides (e.g., hesperetin 7-O-glucoside), and lignan-related molecules. In addition, key thiol-based metabolites involved in redox homeostasis and metal detoxification, such as glutathione (GSH) and phytochelatins (PCs), were identified. The extract also contained osmoprotectants and compatible solutes, including proline, trehalose, sugar alcohols (e.g., D-apiitol), and fructan derivatives (e.g., inulobiose), along with organic acids such as aconitic acid, ribonic acid, and L-tartaric acid. Furthermore, LC-MS analysis further indicated the presence of several phytohormones and their derivatives, including methyl jasmonate, salicylic acid, kinetin, indole-3-butyric acid (IBA), indole-3-acetonitrile (IAN), and auxin conjugates. Lipid-associated molecules, including phosphatidylserine derivatives and sphingolipid precursors, were also identified. Additionally, metabolites associated with photosynthetic and photorespiratory pathways (e.g., phosphoglycolate and hydroxypyruvate), as well as defense-related compounds such as glucosinolates (e.g., sinigrin), phytoalexins, and phenolic esters, were detected, indicating a metabolite profile enriched in stress-responsive bioactive compounds (Table 1; Supplementary Material 2).

**Table 1:**
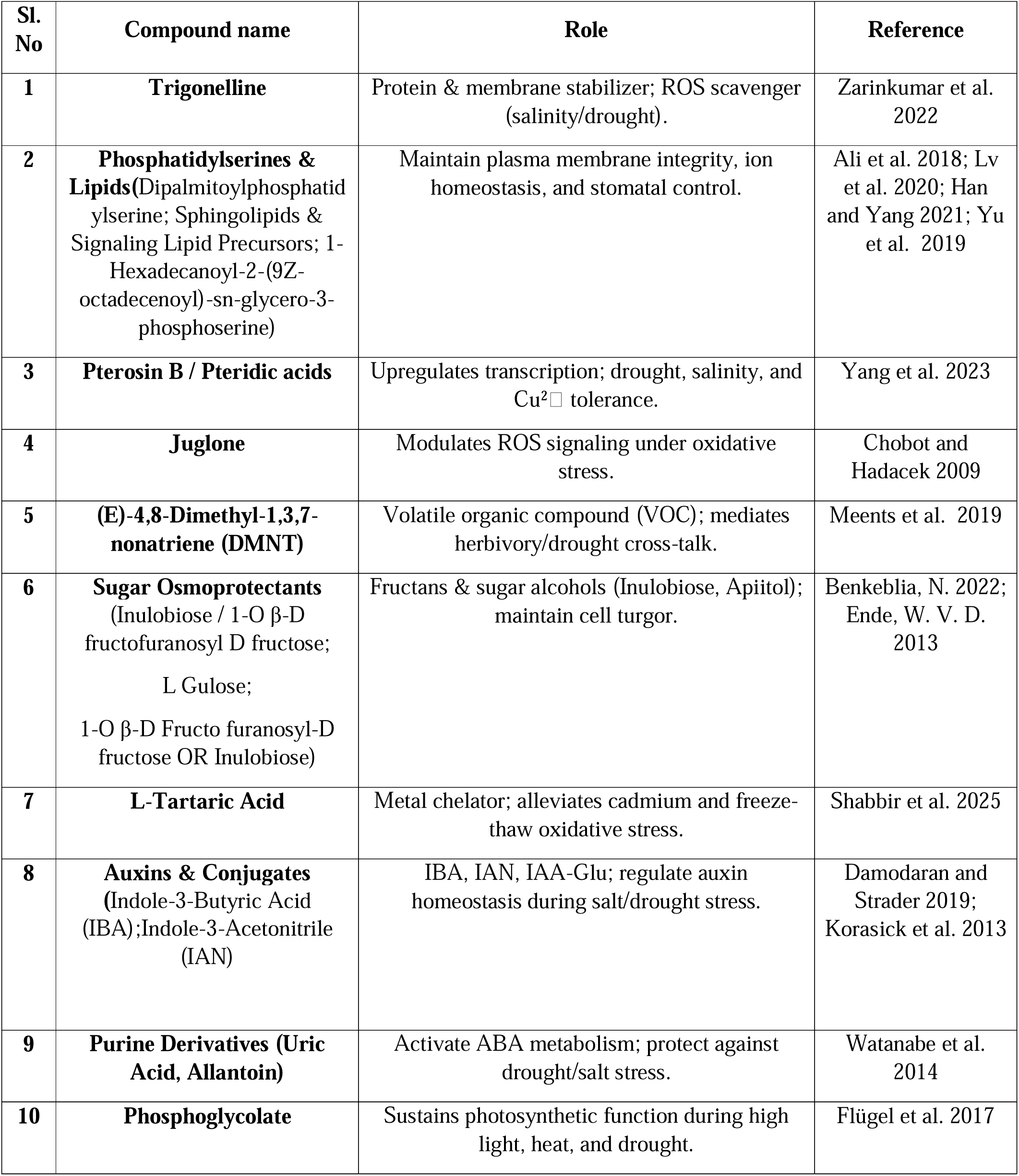
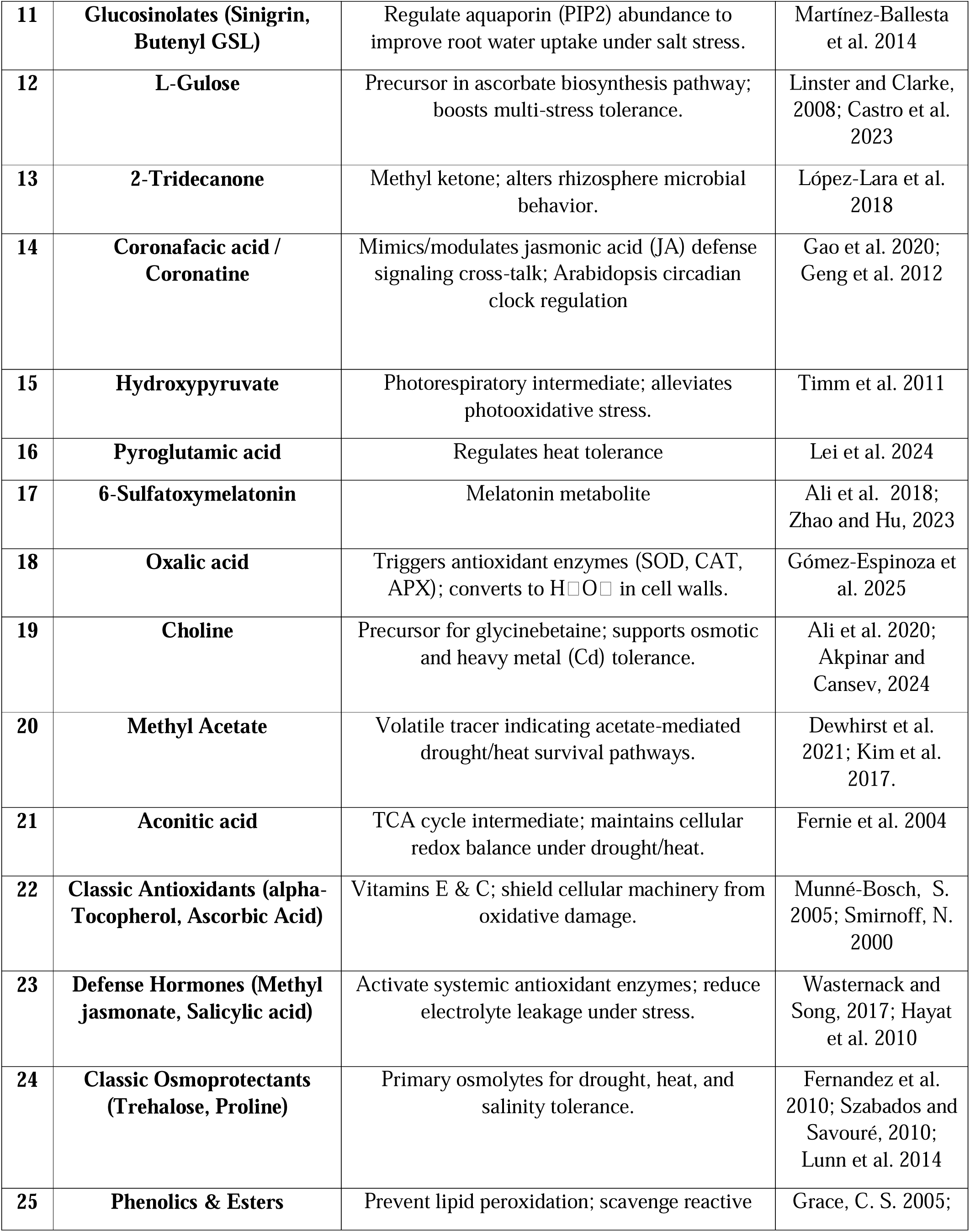
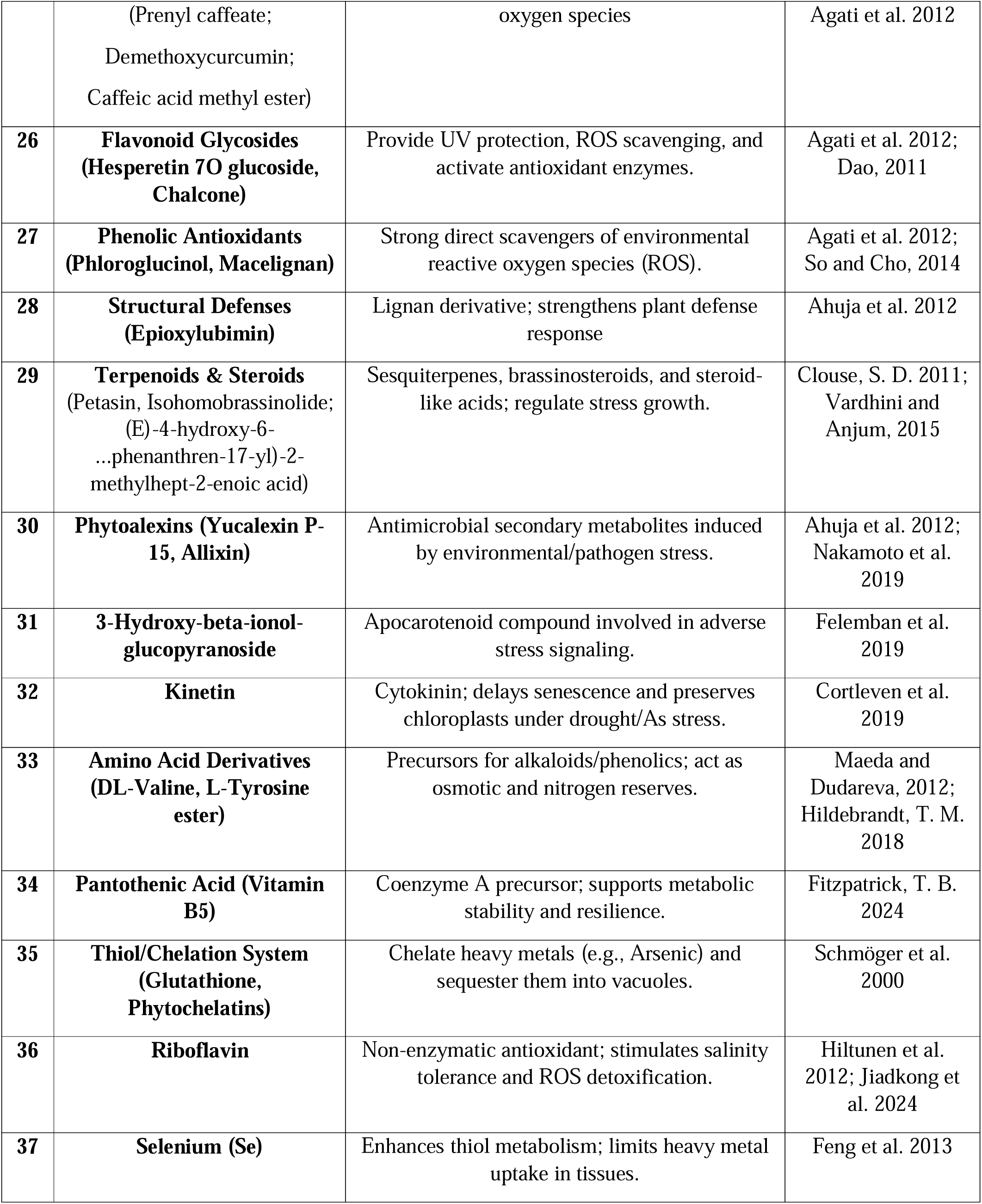
Plant stress-related metabolites identified by LC–MS analysis and their functional roles.

### 3.2 AvE priming restores germination under As stress

Exposure to 10 mg L□¹ As(V) significantly reduced germination in unprimed seeds, with germination frequency declining from 100% (control) to 49.6% and 55.9% at 7 and 14 DAS, respectively (Fig. 1; Table 2). In contrast, AvE priming markedly improved germination under As stress. At 3 DAS, germination increased to ∼65.04% and ∼68.38% in seeds primed with 5 mg mL□¹ and 10 mg mL□¹ AvE, respectively. This recovery was further enhanced at later stages, reaching 82.5% and 74.7% at 7 DAS, and 71% and 75% at 14 DAS for the respective treatments (Fig. 1; Table 2). These results demonstrate the strong ameliorative effect of AvE priming on As-induced inhibition of seed germination. However, when the rice plants were grown under 10 mg L□¹ As(III) stress, no significant difference were observed in physiological parameters in primed plants compared to unprimed plants (Supplementary Material 3)

**Figure 1.**
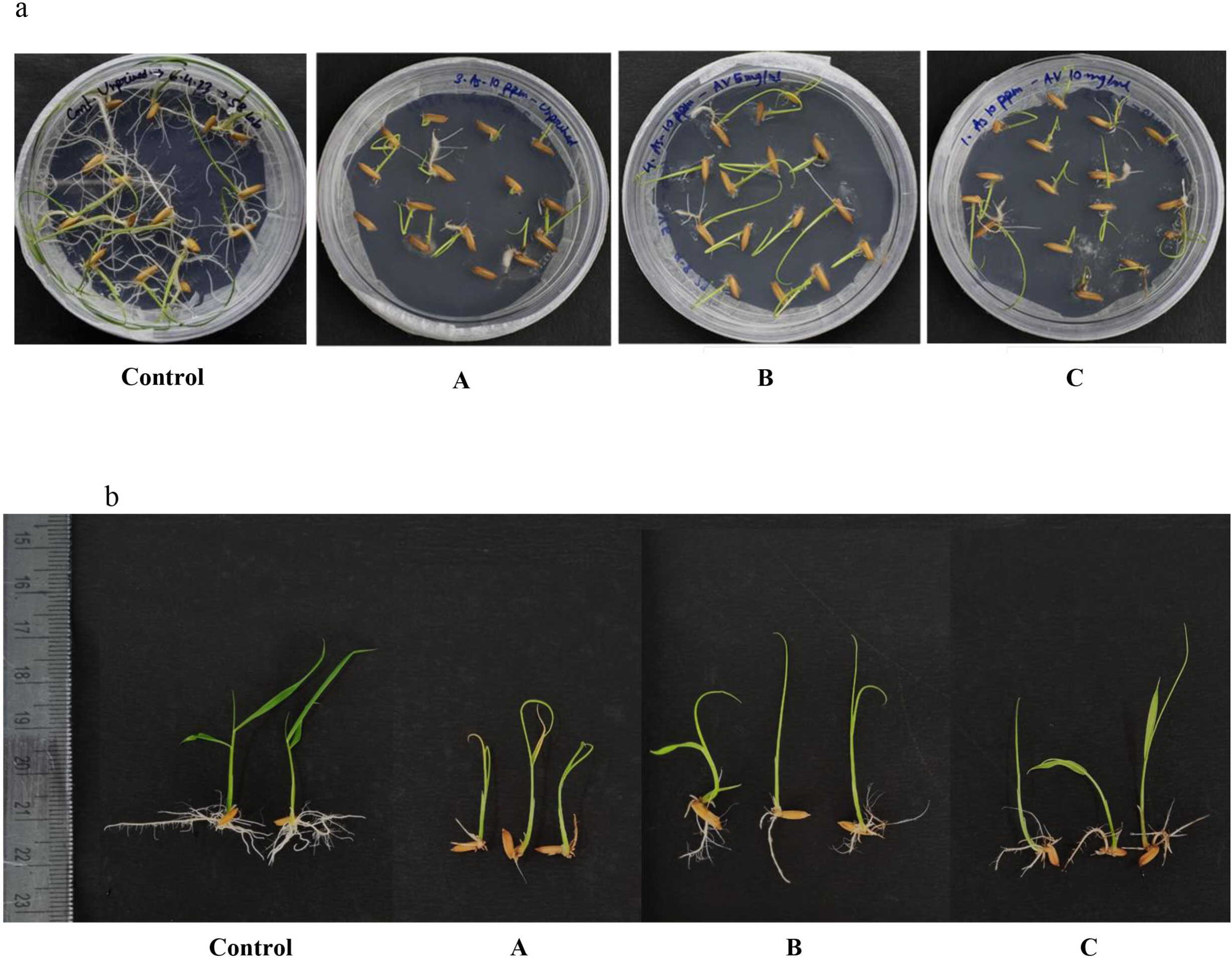
Effects of AvE seed priming on the germination efficiency and seedling growth of rice under As(V). **(a)** Germination of rice seeds grown on water agar plates under (A) As(V) + unprimed, (B) As(V) + AvE 5 mg mL□¹, and (C) As(V) + AvE 10 mg mL□¹ treatments. **(b)** Representative images of 14-day-old rice seedlings (14 DAS) under Control, (A) As(V) + unprimed, (B) As(V) + AvE 5 mg mL□¹, and (C) As(V) + AvE 10 mg mL□¹ treatments.

**Table 2:**
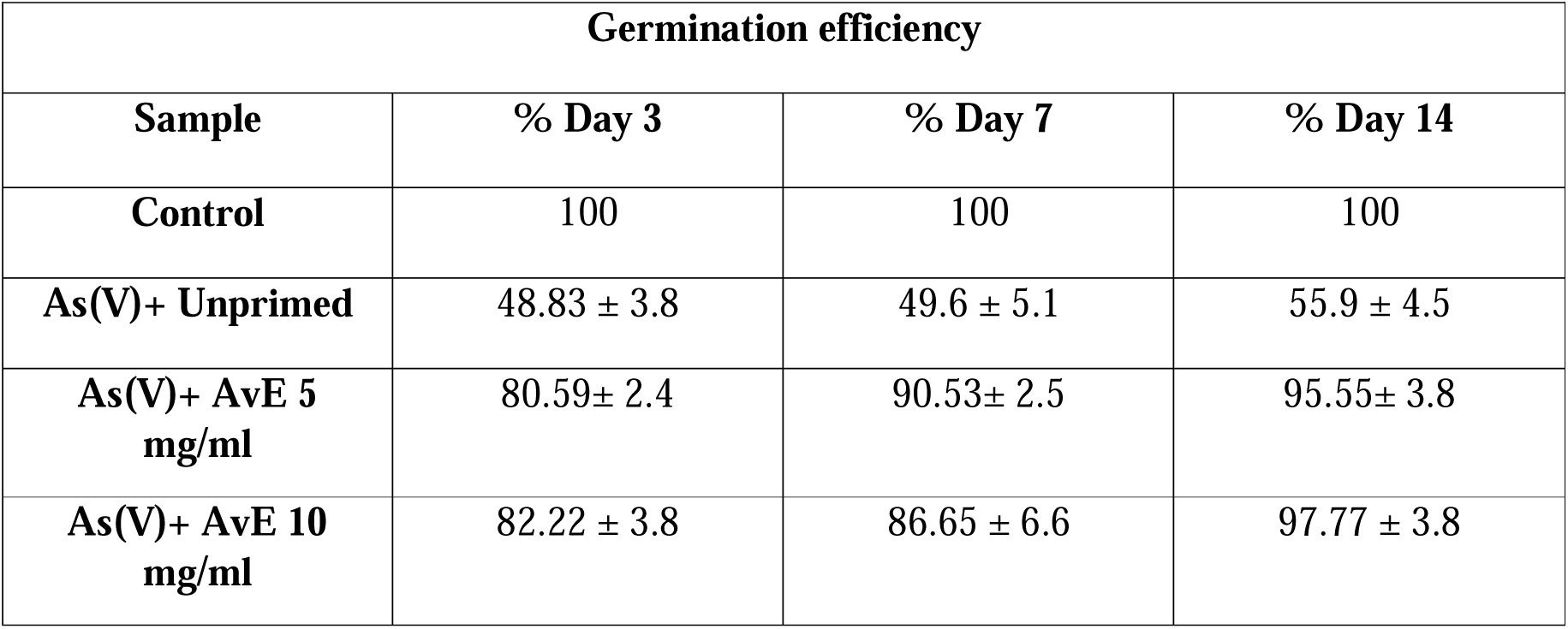
Germination efficiency of rice seeds under Control, As(V) + unprimed, As(V) + AvE 5 mg mL□¹, and As(V) + AvE 10 mg mL□¹ treatments at 3, 7, and 14 days after sowing (DAS).

### 3.3 AvE priming improves seedling growth, biomass, and physiological status under As stress

Arsenic exposure severely impaired seedling growth, as evidenced by reduced root and shoot elongation in unprimed seedlings. However, AvE priming significantly alleviated these effects. Root length increased by ∼1.79–1.99-fold, while shoot length improved by 29–51% in primed seedlings compared to unprimed counterparts under As stress (Fig. 2a–b). Similarly, biomass accumulation was markedly enhanced following priming. Fresh weight and dry weight increased by 1.44–1.74-fold and 1.31–1.45-fold, respectively, relative to unprimed seedlings (Fig. 2c-d). Relative water content (RWC), which declined under As stress, was significantly restored in primed seedlings, showing up to a 1.2-fold increase compared to unprimed plants (Fig. 2e). Chlorophyll content was also adversely affected by As stress; however, AvE priming significantly enhanced chlorophyll a (up to 2.17-fold) and chlorophyll b (68–79%) (Fig. 2f-g). Consistent with these observations, both vigor index I and II were significantly elevated in primed seedlings, showing 1.62–2.01-fold and 3.01– 3.26-fold increases, respectively, compared to unprimed seedlings under As stress (Fig. 2h-i).

**Figure 2.**
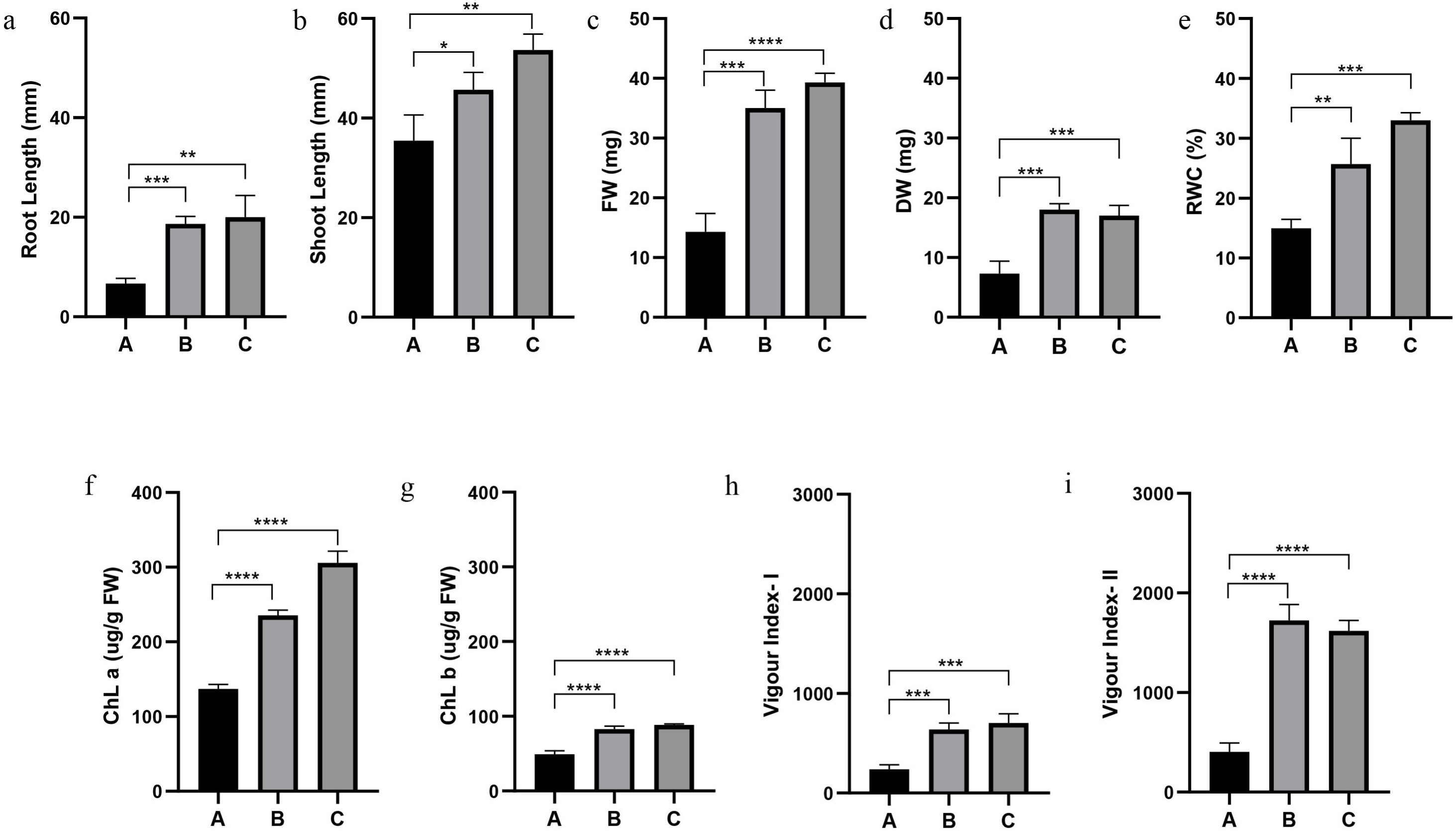
Effects of AvE seed priming on the morphophysiological characteristics of rice seedlings under As(V) stress. The figure shows **(a)** root length, **(b)** shoot length, **(c)** fresh weight (FW), **(d)** dry weight (DW), **(e)** relative water content (RWC), **(f)** chlorophyll *a* (Chl *a*), **(g)** chlorophyll *b* (Chl *b*), **(h)** Vigour Index I, and **(i)** Vigour Index II under (A) As(V) + unprimed, (B) As(V) + AvE 5 mg mL□¹, and (C) As(V) + AvE 10 mg mL□¹ treatments. Data are presented as the mean ± standard deviation (SD) of three biological replicates (n = 3).

### 3.4 AvE priming mitigates oxidative stress and enhances antioxidant defence

Arsenic stress significantly increased oxidative damage, as indicated by elevated levels of H□O□, malondialdehyde (MDA), and proline in unprimed seedlings. H_2_O_2_ levels increased by ∼66.9% in roots and 2.5-fold in shoots under As stress. AvE priming significantly reduced H□O□ accumulation by 23–31% in roots and 21–38% in shoots (Fig. 3a-b). Similarly, MDA content, an indicator of lipid peroxidation, was significantly elevated under As stress but was reduced by 13–18.9% (roots) and 16–26% (shoots) upon AvE priming (Fig. 3c-d). Proline accumulation also increased substantially under stress (4.17-fold in roots and 2.14-fold in shoots), whereas priming reduced proline levels by 18.9–24% in roots and 34-44.7% in shoots (Fig. 3e-f). In contrast, antioxidant metabolites were significantly enhanced in primed seedlings. Total phenolic and flavonoid contents, which declined under As stress in unprimed plants, were restored and significantly elevated in primed seedlings. Phenolic content increased up to 2.34-fold in roots and ∼55% in shoots (Fig. 3g-h), while flavonoids increased up to 2.43 fold in roots and 2.01 fold in shoots (Fig. 3i-j). Glutathione (GSH) content, which decreased by ∼30.8% under As stress, was significantly restored in primed seedlings, showing an increase of up to 41% (Fig. 3k). Enzymatic antioxidant activity was also modulated by AvE priming. Under As stress, SOD and CAT activities declined in unprimed seedlings, whereas priming significantly enhanced SOD (up to 1.29 fold in roots and 3.6-fold in shoots) (Fig. 4a-b) and CAT activity in both roots (up to 2.7 fold) and shoots (up to 21%) (Fig. 4c-d). Conversely, stress-induced elevations in APX and GPX activities were moderated following priming (Fig. 4e-h), indicating restoration of redox balance.

**Figure 3.**
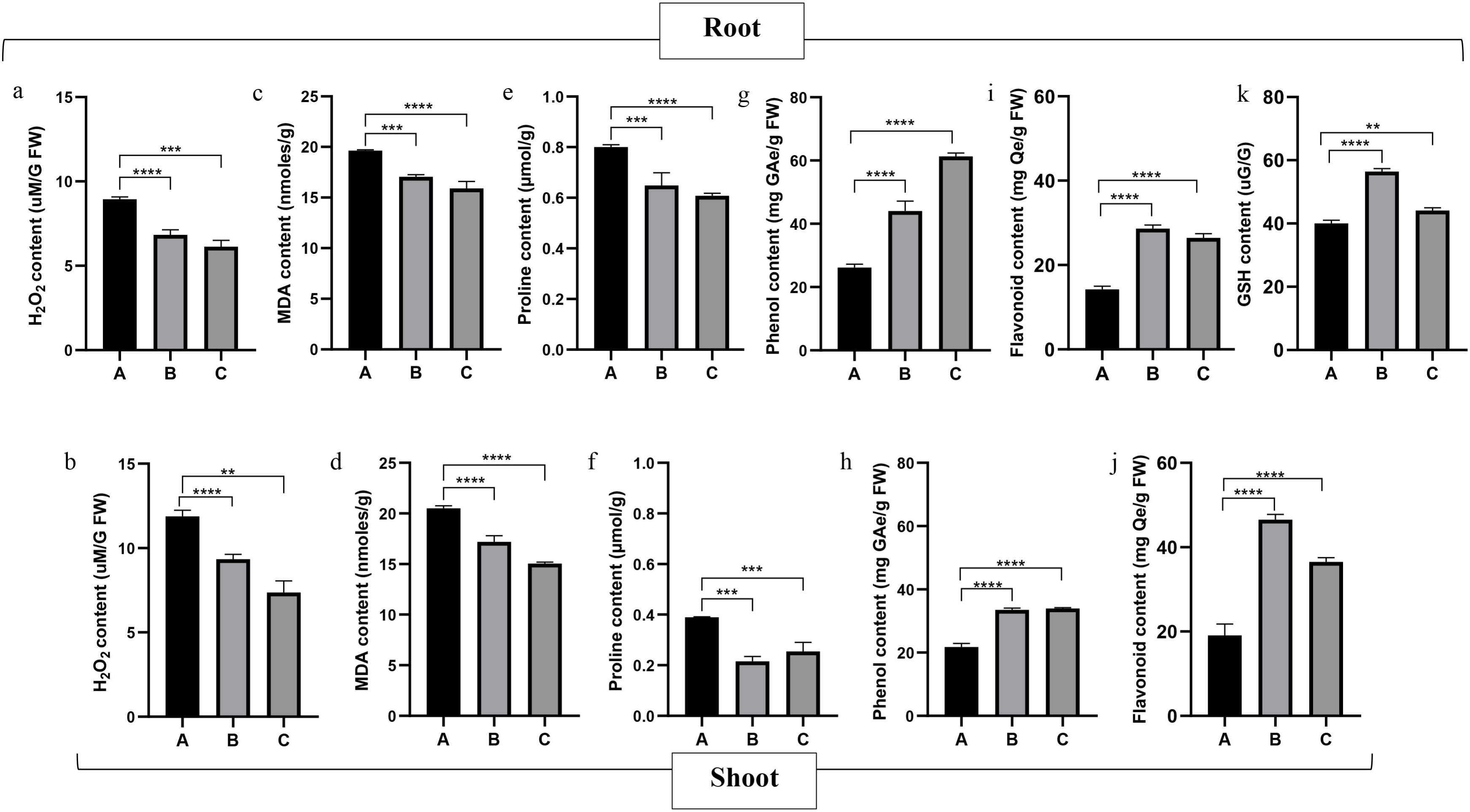
Effect of AvE seed priming on the alleviation of As-induced oxidative stress through the stimulation of antioxidant production in the roots and shoots of 14-day-old (14 DAS) rice seedlings. **Upper panel: (a)** H□O□content, **(c)** TBARS (MDA) content, **(e)** proline content, **(g)** total phenolic content, **(i)** total flavonoid content, and **(k)** GSH content in roots under (A) As(V) + unprimed, (B) As(V) + AvE 5 mg mL□¹, and (C) As(V) + AvE 10 mg mL□¹ treatments. **Lower panel: (b)** H□O□content, **(d)** TBARS (MDA) content, **(f)** proline content, **(h)** total phenolic content, **(j)** total flavonoid content, and **(l)** GSH content in shoots under (A) As(V) + unprimed, (B) As(V) + AvE 5 mg mL□¹, and (C) As(V) + AvE 10 mg mL□¹ treatments. Data are presented as the mean ± standard deviation (SD) of three biological replicates (n = 3).

**Figure 4.**
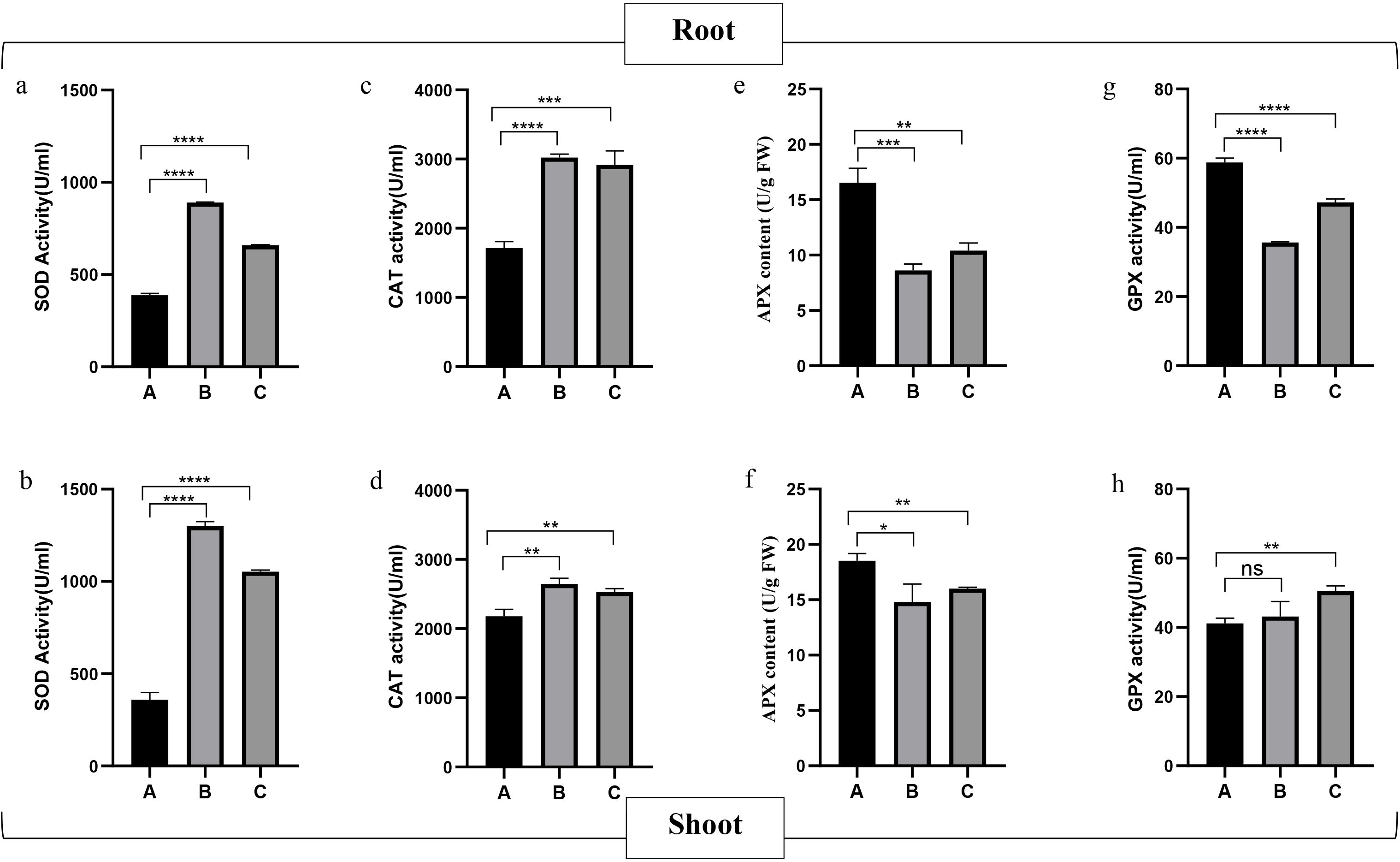
Effect of AvE seed priming on the alleviation of As-induced oxidative stress through the stimulation of antioxidant enzyme activities in the roots and shoots of 14-day-old (14 DAS) rice seedlings. **Upper panel: (a)** SOD activity, **(c)** CAT activity, **(e)** APX activity, and **(g)** GPX activity in roots under (A) As(V) + unprimed, (B) As(V) + AvE 5 mg mL□¹, and (C) As(V) + AvE 10 mg mL□¹ treatments. **Lower panel: (b)** SOD activity, **(d)** CAT activity, **(f)** APX activity, and **(h)** GPX activity in shoots under (A) As(V) + unprimed, (B) As(V) + AvE 5 mg mL□¹, and (C) As(V) + AvE 10 mg mL□¹ treatments. Data are presented as the mean ± standard deviation (SD) of three biological replicates (n = 3).

### 3.5 AvE priming alleviates As-induced anatomical damage

Under As stress, the anatomical structure in roots of rice plantlets displayed significant alterations. A comparative analysis with the control group, where all tissues remained visible and intact, highlighted pronounced distortions, compression of aerenchyma and cortical space, and a reduction in root hairs in unprimed samples subjected to As stress. Notably, in samples primed with AvE, the anatomical structure closely resembled that of the control, with clear visibility of aerenchyma, endodermis, epidermis, xylem, phloem, and turgid root hairs, clearly indicating the ameliorative effect of AvE priming, effectively mitigating As-induced damages to the anatomical structure of both roots and shoots (Fig. 5). To further substantiate the differences in anatomical structure, scanning electron microscopy (SEM) analysis of the root samples was conducted, yielding consistent images that supported the aforementioned findings (Fig. 5). The ultrastructure analysis of rice root transverse sections clearly elucidated the difference in root organisation in the presence of As while AvE priming ameliorated the structural distortion in comparison to unprimed plants. In order to study the structural difference in root anatomy, the average distance between the epidermis and the endodermis of root TS was calculated. The findings depicted that the cortex region of the unprimed plants in the presence of As was highly compressed (-40%) while AvE priming resulted in an increment of the distance between the epidermis and endodermis by 47%-69% in order to ameliorate the effects of As stress in rice root organisation (Table 3). Interestingly, EDAX analysis of these samples indicated the presence of As only in the vascular bundles of the unprimed plants under As stress, but not in the vascular bundles of the AvE-primed plants (Supplementary Material 4).

**Figure 5.**
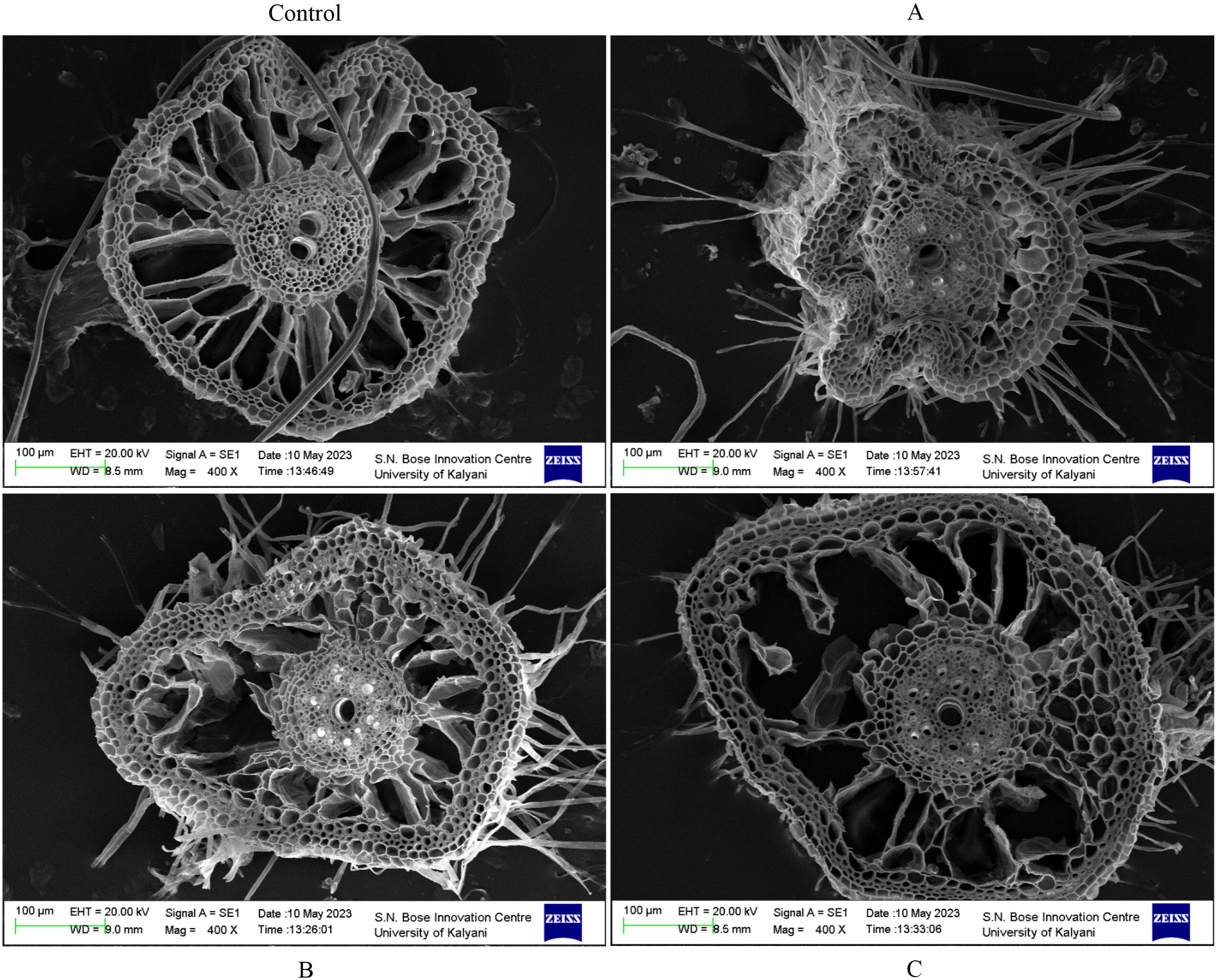
Scanning electron microscopy (SEM) analysis of the root ultrastructure of 14-day-old (14 DAS) rice seedlings under Control, (A) As(V) + unprimed, (B) As(V) + AvE 5 mg mL□¹, and (C) As(V) + AvE 10 mg mL□¹ treatments, demonstrating the anatomical alterations in root architecture.

**Table 3:** Root epidermis-to-endodermis distance in rice seedlings under Control, As(V) + unprimed.

| **Samples** | **Length ($\mu\text{m}$)** |
| --- | --- |
| **Control** | 149.93±0.62 |
| **As(V)+Unprimed** | 89.47±1.68 |
| **As(V)+AvE 5mg/mL** | 131.53±1.96 |
| **As(V)+AvE 10mg/mL** | 151.51±3.3 |

As(V) + AvE 5 mg mL<sup>-1</sup>, and As(V) + AvE 10 mg mL<sup>-1</sup> treatments.

### 3.6 AvE priming reduces arsenic translocation and grain accumulation

ICP–OES analysis revealed substantial accumulation of As in roots and shoots of unprimed seedlings under stress. However, AvE priming significantly reduced As accumulation by 8.5– 39% in roots and 6–74% in shoots (Fig. 6a-b). Furthermore, root-to-shoot translocation of As was markedly reduced in primed plants. The translocation factor decreased from 38.79% in unprimed seedlings to 16.22% and 25.74% in 5 mg mL□¹ and 10 mg mL□¹ AvE treatments, respectively (Table 4). Most importantly, a corresponding reduction in grain As content was also observed, with a significant decrease by ∼13.7-95% in the treated rice plants compared to the unprimed ones (Fig. 6c).

**Figure 6.**
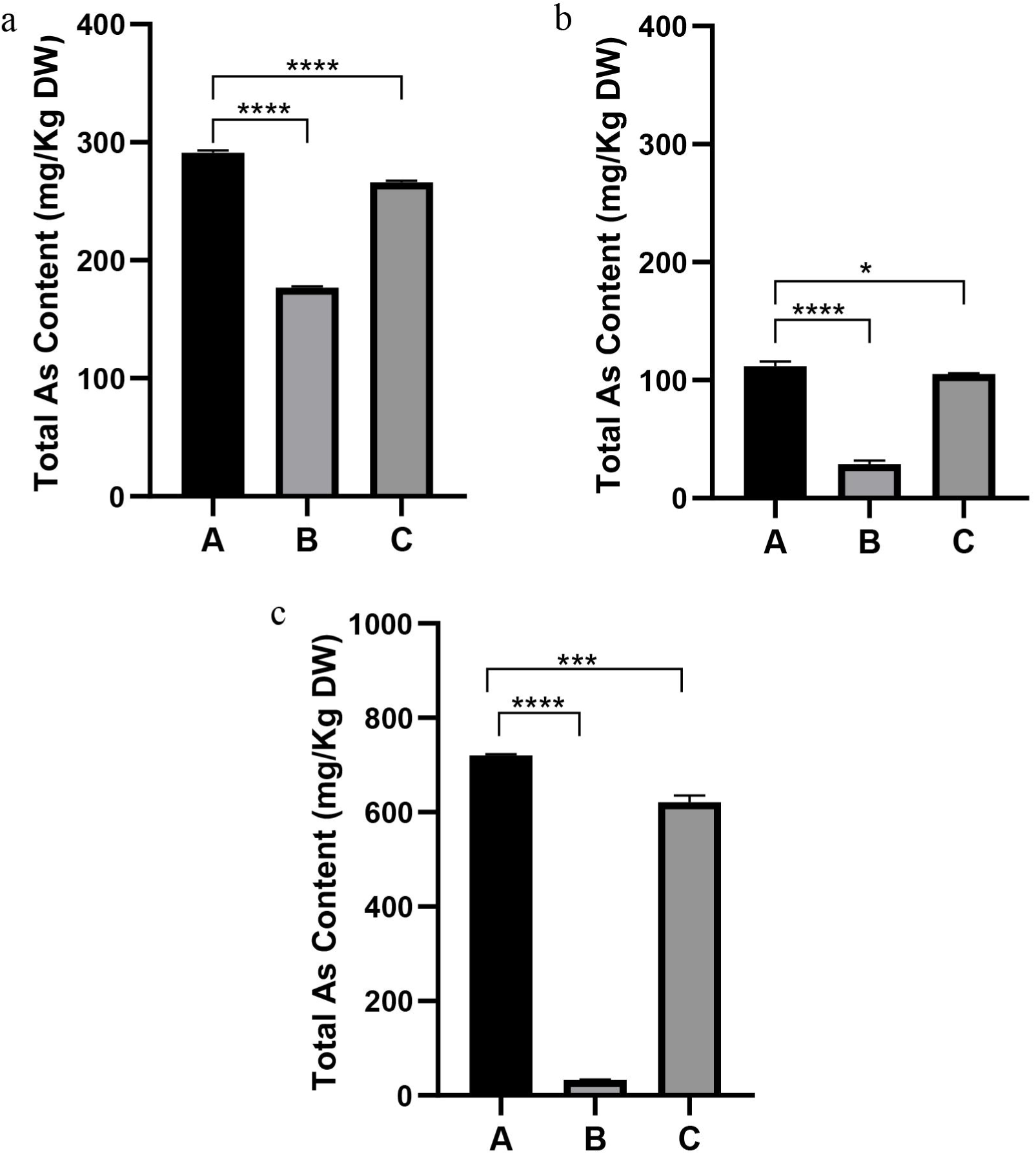
Effects of AvE seed priming on arsenic (As) uptake in (a) roots and (b) shoots of 14-day-old (14 DAS) rice seedlings, and As accumulation in **(c)** grains of mature rice plants under (A) As(V) + unprimed, (B) As(V) + AvE 5 mg mL□¹, and (C) As(V) + AvE 10 mg mL□¹ treatments, as determined by inductively coupled plasma–optical emission spectrometry (ICP-OES).

**Table 4:** Root-to-shoot translocation of arsenic (As) in 14-day-old (14 DAS) rice seedlings.

| Translocation factor (%) |  |
| --- | --- |
| As(V)+Unprimed | 38.79±1.1 |
| As(V)+AvE 5mg/mL | 16.22±1.3 |
| As(V)+AvE 10mg/mL | 25.74±1.1 |

### 3.7 AvE priming modulates expression of stress-responsive genes

To elucidate the effect of AvE priming on the differential expression of morphological (*OsRBOHC*, and *OsSHR2*) and stress (*OsABCC1*, *OsARM1*, *OsHAC1;1*, *OsLSI1*, *OsNRAMP1*, *OsPT1*, *OsPT8*, and *OsCLT1*) marker genes in the root of 14-day seedlings under As stress, the relative expression regulation was quantified using qRT-PCR. Our analysis indicated that, in the roots of unprimed rice plants, the relative expression of *OsABCC1* (1.89 fold), *OsNRAMP1* (20.28 fold), and *OsCLT1* (1.66 fold) showed elevated expression due to exposure to As stress (Supplementary Material 6). On the contrary, the expression of *OsPT1* (0.24 fold), *OsPT8* (0.83 fold), *OsHAC1;1* (0.98 fold), *OsLSI1* (0.04 fold), *OsARM1* (0.71 fold), *OsRBOHC* (0.21 fold), and *OsSHR2* (0.12 fold) was found to be downregulated in presence of As(V) in unprimed roots (Supplementary Material 6). Interestingly, AvE priming was found to have a distinct regulatory role on the expression of the selected genes under As stress. Priming of AvE 5 mg mL^-1^ and AvE 10 mg mL^-1^resulted in the elevated expression of *OsABCC1* (log2FC 0.24 and 0.47, respectively), *OsCLT1* (log2FC 0.77 and 0.09, respectively), *OsHAC1;1* (log2FC 0.62 and 1.12, respectively), *OsPT8* (log2FC 0.7 and 0.14, respectively), and *OsSHR2* (log2FC 1.7 and 1.48, respectively) relative to unprimed roots (Fig. 7). However, the regulatory impact on other key morphological and stress markers was highly concentration-dependent. AvE 5 mg mL^-1^ was proven highly effective in upregulating stress markers such as *OsARM1* (log2FC 0.37), *OsLSI1* (log2FC 2.34), *OsNRAMP1* (log2FC 0.27), and *OsPT1* (log2FC 0.80) along with morphological marker *OsRBOHC* (log2FC 1.23). Contrary to that, higher concentration AvE 10 mg mL^-1^ was found to be resisting the translational expression of *OsARM1* (log2FC -0.84), *OsLSI1* (log2FC -0.62), *OsNRAMP1* (log2FC -0.89), *OsPT1* (log2FC -0.85), and *OsRBOHC* (log2FC -0.40) (Fig. 7).

**Figure 7.**
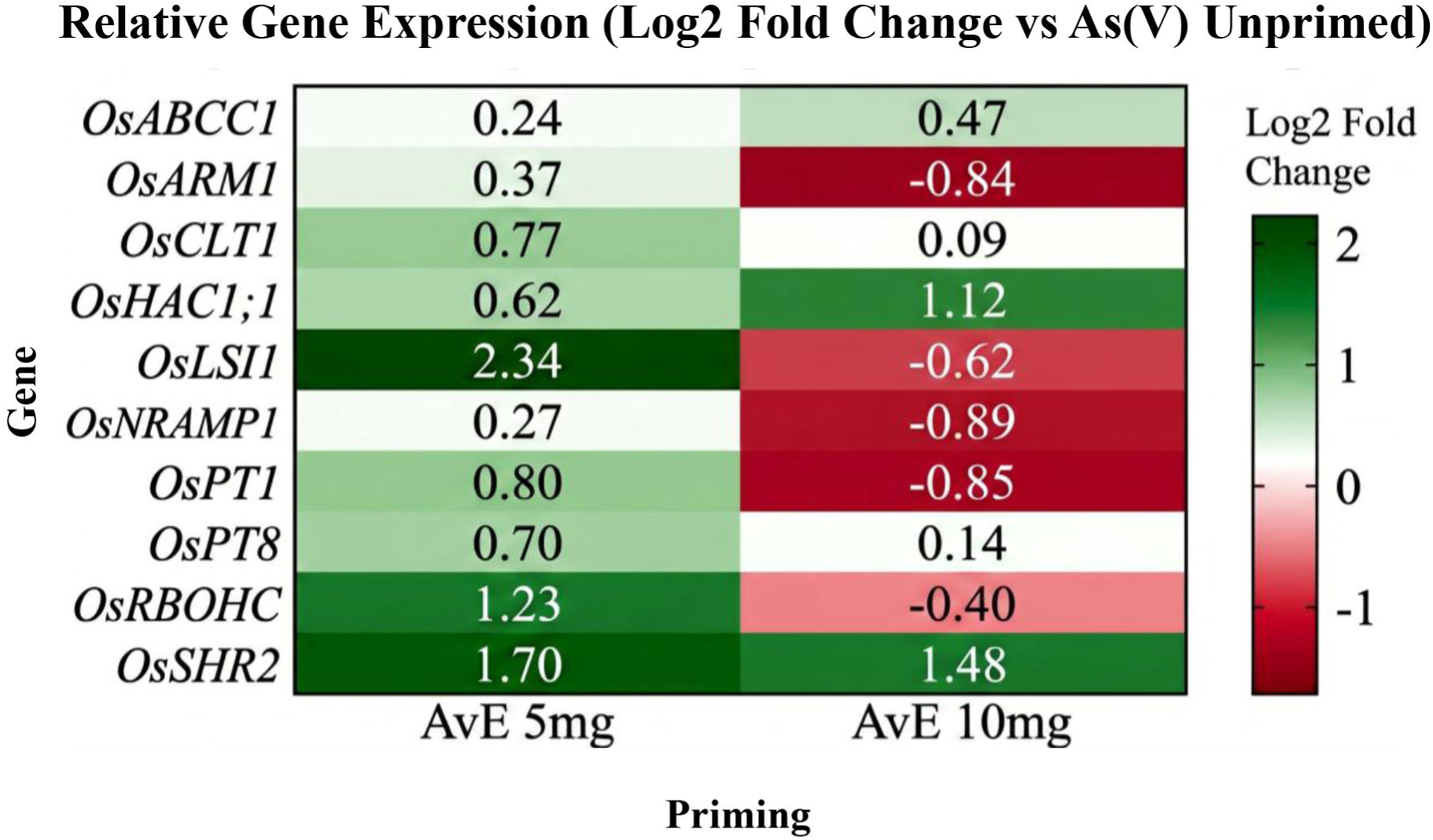
Quantitative real-time PCR (qRT-PCR)-based relative expression analysis of stress-responsive genes (*OsABCC1, OsARM1, OsCLT1, OsHAC1;1, OsLSI1, OsNRAMP1, OsPT1*, and *OsPT8*) and root morphological marker genes (*OsRBOHC* and *OsSHR2*) in the roots of 14-day-old (14 DAS) rice seedlings under As stress following dose-dependent AvE seed priming. *OsHisH3* was used as the endogenous control.

### 3.8 AvE priming enhances agronomic performance

Field evaluation revealed significant improvements in agronomic traits in AvE-primed plants under arsenic stress. Primed plants exhibited earlier flowering (up to 7 to 10 days), increased plant height (13.5–28%), and higher tiller numbers (52.7–68.5%). Yield-associated traits, including panicle number (up to 1.9 folds), panicle length (47.8–58%), and grain number per panicle (50–75%) were significantly improved. Grain filling percentage was increased by 2.3–4.2 folds in primed treatments, demonstrating the translational potential of AvE priming in enhancing crop productivity under stress conditions (Table 5; Fig. 8a,b).

**Table 5:** Agronomic parameters of mature rice plants under As(V) + unprimed, As(V) + AvE 5 mg mL□¹, and As(V) + AvE 10 mg mL□¹ treatments.

| No. | Agronomic Trait | As(V)+Unprimed | As(V)+AvE 5<br>mg/mL | As(V)+AvE 10<br>mg/mL |
| --- | --- | --- | --- | --- |
| 1. | Plant height (cm) | 89±2.20 | 114±0.90 | 101±1.34 |
| 2. | Tiller Number | 6.33±0.58 | 10.67±0.58 | 9.67±0.58 |
| 3. | Panicle Number | 4.67±0.58 | 13.67±2.08 | 12±2 |
| 4. | Panicle Length (cm) | 15±0.5 | 23.67±0.76 | 22.17±0.76 |
| 5. | DAS to Reproductive Maturity (days) | 97± 7 | 87± 4 | 90±4 |
| 6. | 25 Grain Weight (mg) | 410±1.15 | 478±1 | 420±2 |
| 7. | Grain Length (mm) | 8.37±0.48 | 10±0.5 | 8.625±0.48 |
| 8. | Grain Width (mm) | 2±0 | 2.4±0.48 | 2.1±0.33 |
| 9. | Number of filled grain | 22±4 | 92±3 | 52±2 |
| 10. | Grain per panicle | 4±0.25 | 7±0.33 | 6±0.25 |

**Figure 8.**
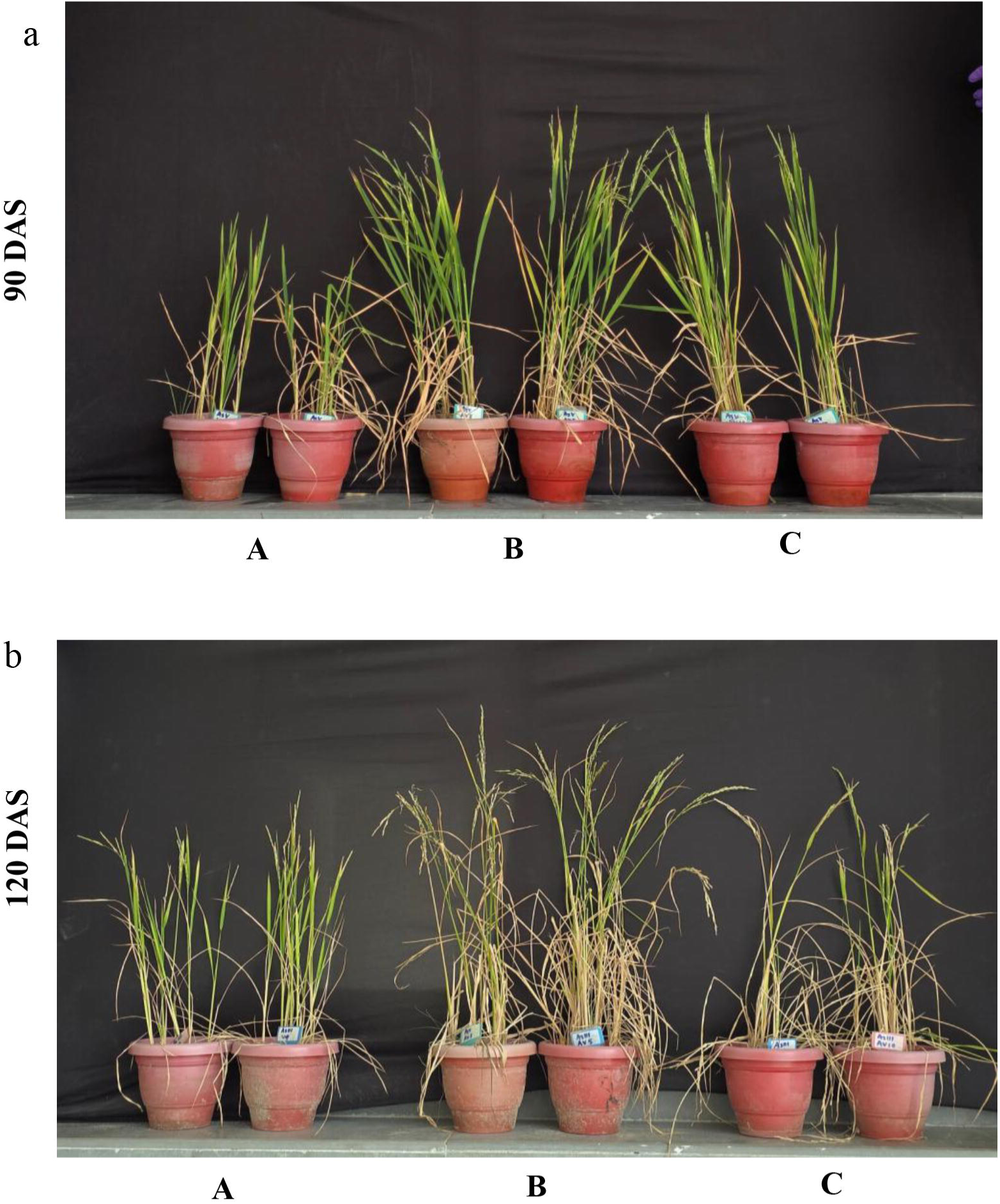
Effect of AvE seed priming on the growth of mature rice plants under As stress, showing the phenotypic appearance at **(a)** 90 DAS and **(b)** 120 DAS under (A) As(V) + unprimed, (B) As(V) + AvE 5 mg mL□¹, and (C) As(V) + AvE 10 mg mL□¹ treatments.

## 4. Discussions

Groundwater contamination by As in the Gangetic alluvial zones of West Bengal is estimated to span approximately 39,000 square kilometres and has evolved into a significant health concern related to As exposure, affecting 42.7 million people across 85 blocks. Historically, the primary focus has been on As contamination in drinking water sourced from groundwater. However, a recent shift in attention has acknowledged the agricultural sector, which heavily relies (>90%) on the same contaminated groundwater for irrigation. The consumption of these As-laden foodstuffs, in addition to the intake of As-contaminated drinking water, has significantly worsened the As-poisoning scenario in the Bengal Basin. Additionally, this spread of As via agricultural produce, especially rice, has significantly multiplied the threat of As toxicity, resulting in As-related health hazards even in the areas not affected by As-contaminated groundwater. As a result, the Bengal Basin, a component of the larger Ganga-Brahmaputra-Meghna river basin and inhabited by millions of people, has witnessed the highest incidence of individuals impacted by As toxicity (Das et al. 2021, Sarkar and Paul 2016, Smits et al. 2019).

To reduce As accumulation in rice grains, agronomic practices, bioremediation, or application of molecular biology techniques have been employed (Das et al. 2020). However, the cost involved in controlling As in soils has made genetic engineering the strategy of choice to curb As accumulation in rice grains in modern times. Nevertheless, the fact that the toxic metalloid As uptake and transport in rice is mediated by endogenous transporters due to the imperfect selectivity while acquiring necessary elemental ions from the soil (Das et al. 2020) makes developing an efficient biotechnological solution one of the biggest challenges for mankind. Still, considering the serious threat posed by As-accumulation in rice grains, several genetic engineering approaches have been undertaken to minimise As-accumulation in rice grains with limited success. Although plant genetic modification has been widely used for improving As-stress mitigation in rice plants, the genetic engineering process is associated with several constraints, including regulatory, ecological, and societal constraints, limiting their widespread adoption. Therefore, the exploration of alternative methods to genetic modification for enhanced As-stress resistance has become popular in recent times. Interestingly, seed priming technology that involves the immersion of seeds in water or a solution containing various substances, followed by drying, either before or after sowing in beds or directly in the field, has become a modern tool to mitigate abiotic stress in plants (Harris et al. 2008). Positive effects of seed priming have been observed across diverse abiotic stress conditions in different crops. Seed priming has demonstrated efficacy in alleviating cold and salt stress in capsicum, as well as mitigating salinity and drought stress in sugarcane and mustard (Yadav et al. 2011, Patade et al. 2009). Additionally, favorable outcomes of seed priming with selenium solution have been documented in various field crops and vegetables (Hasanuzzaman et al. 2020, Chen & Sung, 2001). Further, the utilization of nanoparticles as priming solution has also emerged as a popular and promising approach to mitigate the toxicity of heavy metals and metalloids in plants. Zinc Oxide Nanoparticles (ZnONPs) have gained widespread application as nano fertilizers, particularly for addressing zinc deficiency in agricultural soils (Ma et al. 2020). ZnONPs have been demonstrated to reduce the accumulation of As, concurrently promoting growth and photosynthesis in rice seedlings (Yan et al. 2021). Similarly, the plant growth promoting microelement potassium humate upon priming of rice seeds significantly improved seed germination, seedling growth, and antioxidant defense system under As stress, further establishing the efficiency of seed priming as an emerging dynamic approach for regulating the seed germination and plant growth under As stress conditions (Mridha et al. 2021). Since, seed priming is estimated to promote elevated seed emergence rate, uniform germination, and robust seedling growth that are directly impeded by As, employing seed priming stands out as a viable strategy to rapidly enhance both the germination rate and early growth by modulating physiological parameters (Acharya et al. 2020). Although a few studies have been performed concerning seed priming to improve As stress mitigation in rice, so far there is no report regarding the use of phyto-extracts from terrestrial weeds-mediated seed priming in reducing the As toxicity in rice. The phytoextracts of terrestrial weeds are a rich source of antioxidants, phytohormones, and mineral nutrients that have the potency to act as a natural bio-stimulant. However, the application of these phytoextracts from terrestrial weed-mediated seed priming in regulating the As toxicity in rice seedlings has not been studied till date. Therefore, the current study proposes a novel non-transgenic seed priming method that employs phytoextracts derived from agricultural weeds to mitigate As-toxicity and reduce As accumulation in grains of rice plants cultivated in As-contaminated areas. The hypothesis is that this innovative phytopriming approach will activate signalling pathways, leading to faster defense responses to As stress-induced oxidative damages, resulting in the development of a more resilient rice variety with reduced accumulation of As. The proposed study, potentially the first study of its kind, aims to (i) examine whether priming rice seeds with terrestrial weed phytoextracts enhances seed germination and seedling growth in comparison to unprimed seeds, (ii) assess the levels of stress markers and antioxidant activities in seedlings subjected to priming versus those from unprimed seeds, (iii) compare the morpho-anatomical features between the primed and unprimed rice plantlets, and (iv) compare the accumulation and translocation of As in seedlings subjected to priming and those from unprimed seeds under As-stressed conditions.

In this study, we demonstrate that seed priming with agricultural weed, *Amaranthus viridis* extract (AvE) confers significant tolerance to As stress through a multi-layered mechanism involving redox regulation, osmotic adjustment, metal detoxification, and transcriptional reprogramming. The initial LC–MS profiling revealed that AvE is enriched with diverse bioactive metabolites, including antioxidants (ascorbate, α-tocopherol, phenolics, flavonoids), thiol-based detoxifiers (glutathione and phytochelatins), osmoprotectants (proline, trehalose), phytohormones (jasmonates, salicylates, auxins, cytokinins), and lipid-associated signaling molecules. The coexistence of these metabolite classes suggests that AvE functions as a multi-component biostimulant capable of simultaneously targeting multiple stress-responsive pathways (Table 1).

One of the earliest and most vulnerable stages affected by As stress is seed germination and early seedling establishment. Arsenic interferes with sulfhydryl groups, induces protein carbonylation, and disrupts cellular metabolism, resulting in impaired germination and stunted growth (Chowardhara et al. 2019; Banerjee et al. 2023). In agreement, our study observed a substantial decline in germination under As stress. However, compared to unprimed plants AvE priming was found to significantly improve germination frequency by up to 82.5% and 75% at 7 DAS and 14 DAS, respectively; indicating the positive influence AvE priming during the germination phase of rice plants during As stress (Fig. 1; Table 2). Similarly, AvE priming demonstrated significant efficacy in mitigating As-induced toxicity, resulting in a notable enhancement of root and shoot growth by 1.99-fold and 51%, respectively, when compared to unprimed conditions (Fig. 2a-b). This positive influence on root and shoot development, in turn, led to a substantial increase in rice biomass. AvE priming exhibited concentration-dependent effects on rice fresh weight, demonstrating a remarkable 1.74-fold increment and a 1.45-fold increase in dry weight compared to unprimed under As stress conditions (Fig. 2c-d). Additionally, our study investigated the impact of AvE priming on the water retention capacity (RWC) of rice plants, a parameter indicative of cell membrane integrity under As stress. Previous research has documented a reduction in RWC as a manifestation of cell membrane damage under As stress. In our study, we observed a severe decline in RWC under As stress, consistent with existing literature. Intriguingly, AvE priming demonstrated significant ameliorative effects, elevating the RWC of primed plantlets by 71% and 1.2-fold, respectively, in the presence of As in the growth medium (Fig. 2e). Further, upon entering the plant cells, As significantly affects the availability of Mg ions. Along with playing a critical role as co-factors in various enzymatic activities, Mg ions carry out essential functions in the light-harvesting complexes of chlorophylls. Thus, exposure to As stress alters the light-harvesting complexes and chlorophyll biosynthesis, resulting in the disruption of photosynthetic activities (Emamverdian et al. 2015, Srivastava and Sharma 2014). Consistent with these observations, a drastic reduction in the content of photosynthetic pigments chlorophyll a and chlorophyll b was observed in unprimed seedlings under As stress. Remarkably, prophylactic treatment of AvE was found to increase the amount of chlorophyll a and chlorophyll b by 1.17 folds and 79% respectively under As stress clearly emphasizing the broad applicability of AvE priming in overcoming early-stage stress constraints of rice plants under As-induced stress conditions (Fig. 2f-g).

The central mechanism underlying As toxicity is mediated by excessive generation of reactive oxygen species (ROS) that in turn lead to interference with various cellular processes, particularly in mitochondria and chloroplasts causing an imbalance between ROS production and the plant’s antioxidant defense mechanisms ultimately resulting in oxidative stress damages (Srivastava et al. 2017, Hu et al. 2020). The oxidative stress triggers the oxidation of lipids by removing electrons from the hydrogen atoms within the fatty acyl chains of polyunsaturated fatty acids present in both the cellular and organelle-bound plasma membranes. This process culminates in the production of malondialdehyde (MDA), which causes heightened membrane rigidity, increased leakiness, and damaged membrane proteins. Subsequently, this damage prompts the release of vital cellular components, such as electrolytes, from the cell, ultimately leading to cellular demise (Abbas et al. 2018). Therefore, quantification of hydrogen peroxide (H_2_O_2_) and thiobarbituric acid reactive substances (TBARS, indicative of membrane lipid peroxidation) has been widely used to gauge the severity of membrane damage and ROS accumulation as a measure to assess the As-induced oxidative stress in plant cells (Chowardhara et al. 2022, Banerjee et al. 2023). Interestingly, in our study, the application of AvE priming revealed a noteworthy decrease of H_2_O_2_ (up to 31% in roots and 38% in shoots) (Fig. 3a-b) and TBARS content (up to 18.9% in roots and 26% in shoots) (Fig. 3c-d) in 14 DAS seedlings under As stress, clearly indicating the involvement of AvE priming in increased cell stability under As stress. In addition to H_2_O_2_ and TBARS, proline also serves as a crucial marker for determining cellular oxidative stress, contributing to the regulation of osmolyte homeostasis. An optimal proline concentration plays a pivotal role in governing protein synthesis, maintaining enzyme stability, and acting as a scavenger for hydroxyl radicals. However, surpassing the physiological threshold of proline may exert adverse effects by impeding certain metabolic activities, notably substrate-level phosphorylation (Schat et al., 1997; Bakry et al., 2014). In the present study, AvE priming demonstrated efficacy in mitigating As stress in 14-day-old seedlings, resulting in a noteworthy reduction of proline content in both root (18.9%-24%) and shoot (34%-44.7%) tissues compared to unprimed plants (Fig. 3e-f). The diminished proline levels further underscore the ameliorative impact of AvE priming on the regulatory mechanisms associated with oxidative stress, highlighting its potential in maintaining a balanced cellular environment under As-induced physiological challenges.

Interestingly, priming of rice seeds with AvE significantly augmented the total phenolic content in the root (68% and 2.34-fold, respectively) and shoot (54% and 56%, respectively) under As stress conditions (Fig. 3g-h). Among various cellular antioxidants, phenolics stand out as well-established osmoregulators and antioxidants, contributing to enhanced adaptability under As-induced oxidative stress (Moulick et al. 2016). The results also showed significant increase in flavonoids in primed seedlings in roots (up to 0.9-2.43 folds respectively) and shoots (up to 0.8 - 2.01 folds respectively) (Fig. 3i-j). As plant secondary metabolites, flavonoids play an important role under As stress to mitigate the stress condition by acting as not only antioxidants, heavy metal detoxifiers, and regulators of plant hormones like auxin, but also significantly enhance the plant ability to cope with oxidative stress and the heavy metal toxicity (Sharma et al. 2019; Mathur et al. 2022; Wang et al. 2019). In addition to phenolics and flavonoids, glutathione (GSH) plays a pivotal role in mitigating stress induced by heavy metalloids through chelation mechanisms (Anjum et al. 2016). In our study, a significant reduction in GSH levels in the root of unprimed seedlings (-30.8%) was observed after 14 days of As exposure. This decline may be attributed to the utilisation of GSH in the conversion of As(V) to As(III) during As(V) uptake by the root, coupled with the inhibition of glutathione reductase activity, consequently impeding the GSH recycling process. Conversely, priming with AvE at 5 mg mL^-1^ significantly elevated GSH content in the root (41%), indicating a potential activation of glutathione reductase in stress mitigation (Fig. 3k). While the treatment with AvE at 10 mg mL^-1^ resulted in a slight increase in GSH in the root (10%), the findings suggest a nuanced modulation of glutathione reductase activity in response to AvE priming under As-induced stress conditions (Fig 3E).

The analysis of the enzymatic antioxidant system further corroborates this improved redox balance. Under prolonged As stress, antioxidant enzymes such as SOD and CAT are often inhibited due to oxidative damage, while APX and GPX may be overactivated (Shri et al. 2009; Hasanuzzaman et al. 2020). In the present study, AvE priming improved SOD in both roots (up to 0.69-2.29 folds) and shoots (2.92-3.6 folds) (Fig. 4a-b) and CAT activity in both roots (up to 0.69-2.76 folds respectively) and in shoots (16.25-21.37%) (Fig. 4c-d), compared to unprimed plants under As stress. Most importantly, AvE priming point to a coordinated regulation for sustained stress tolerance by normalising APX in both roots (37-47.95%) and shoot (13.6-20%) (Fig. 4e-f), and GPX activity in shoot (increased by 4.86-22.8%) and root (decreased by 19.63-39.38%) (Fig. 4g-h) respectively, indicating a shift from stress-induced overcompensation to balanced redox homeostasis (Srivastava et al. 2016).

The ameliorative potential of AvE priming countering As stress was further visualized while comparing the anatomical structure of the primed and unprimed seedlings. Generally, the anatomical integrity of the root is profoundly impacted by As stress, resulting in the distortion of both the epidermis and endodermis layers. This perturbation often extends to the vascular bundles responsible for As transport, exhibiting significant structural anomalies. Additionally, As-induced osmotic imbalance and diminished membrane stability contribute to the pronounced distortion of the tissue structure under As stress conditions. In our investigation, the unprimed root anatomy demonstrated noteworthy alterations, especially severe compaction of the cortical tissues due to As stress (Fig. 5), possibly driven by lipid peroxidation and osmotic imbalance (Khan et al. 2021). However, the application of AvE was able to induce a counteractive response, where the roots of primed plants resembled the anatomical structures of plants grown under non-stress conditions (Fig. 5). Moreover, comparative analysis of the root ultrastructure revealed intricately developed aerenchyma tissue in the roots of AvE-primed seedlings even under As stress; whereas in case of the unprimed plantlets exposed to As stress the aerenchyma was found to be poorly developed or absent (Fig. 5). The aerenchyma is a specialized tissue characterized by the presence of air spaces or channels within the root structure and is assumed to play a multifaceted role in mitigating the impacts of As stress. It enhances oxygen supply, reduces hypoxia, limits As uptake, and contributes to overall stress tolerance, thereby aiding in the plant’s adaptation to As-contaminated environments (Ejiri et al. 2021, Mei et al. 2012). Thus, AvE priming is presumed to have a role in the amelioration of anatomical changes, effectively overcoming the distorted root structure and restoring control over the anatomical organisation, thereby mitigating As toxicity in primed rice plantlets.

The most important finding of this study is the significant reduction in As accumulation and root-to-shoot translocation in AvE-primed plants. Relative to unprimed counterparts, AvE priming resulted in a substantial decrease in As uptake by the roots and the overall As content (Fig 6). Furthermore, the study demonstrated that As translocation from root to shoot was significantly minimised, with the translocation factor reduced by up to 58% (Table 4). This marked reduction in the translocation factor in AvE-primed seedlings suggested the activation of specific stress mitigation mechanisms to impede the transport of As from roots to shoots. Also the As content of the mature grain showed 95% downregulation in AvE primed samples (Fig. 6c). Thereafter to unravel the potential molecular mechanisms underlying the alleviation of As stress by AvE priming, we conducted qRT-PCR-based relative expression analyses of select critical genes (Fig 7). The selection of gene candidates was made based on their recognized roles in morphological development and As stress response. Intriguingly, the expression profiles of these genes exhibited a strong correlation with changes in oxidative stress and antioxidant balance, morpho-anatomical structural variations, and the pattern of As accumulation in 14 DAS AvE-primed seedlings compared to their unprimed counterparts. Among the As-responsive genes investigated, OsABCC1, a C-type ATP-binding cassette (ABC) transporter localized in the tonoplast, plays a pivotal role in sequestering AsIII-phytochelatin complexes inside the vacuole of vascular bundles, primarily in phloem companion cells. Previous studies have reported that the overexpression of *OsABCC1* in rice effectively limits As transport to the grain from the root via the shoot by sequestering As into the vacuole of phloem companion cells (Song et al. 2014). Interestingly, our study revealed that exposure to As in unprimed plants led to a slight upregulation of *OsABCC1*, similar to findings by Song et al. (2014) (Fig 7). However, in AvE-primed seedlings, the expression of *OsABCC1* was significantly enhanced, suggesting an elevated sequestration of As in the root (Fig 7). This, in turn, may be responsible for the observed reduction in the root-to-shoot translocation factor in our study. On the other hand, the CRT-like plastid-bound transporter (*OsCLT1*) is known to play a crucial role in maintaining glutathione homeostasis in rice roots. Glutathione, in turn, regulates As-induced cellular oxidative stress by forming phytochelatins that complex with heavy metal(loid) ions and are ultimately sequestered in vacuoles. Previous studies involving *OsCLT1* mutants demonstrated not only reduced accumulation of root As levels but also lower GSH levels in roots, emphasizing the significance of *OsCLT1* in mitigating As stress in rice roots (Yang et al. 2016). Remarkably, AvE priming resulted in a significant upregulation of *OsCLT1* expression in the presence of As, aligning with the increased GSH content in primed roots (Fig 7). Additionally, AvE priming further regulated the expression of several key molecular markers involved in arsenic uptake and detoxification, indicating multi-faceted involvement in the rice As response network. Notably, the enhanced expression of *OsHAC1;1*, the key arsenate reductase enzyme responsible for the conversion of As(V) to As(III), indicating increased intercellular detoxification capacity prior to PC mediated sequestration. Similar to that, the concentration-dependent regulation of *OsPT1*, *OsPT8, OsLSI1*, and *OsNRAMP1* suggests the involvement of AvE priming in arsenic transport and nutrient homeostasis rather than merely arsenic uptake suppression. Furthermore, the AvE priming with 5 mg mL^-1^ was found to have a distinct regulatory role on the relative expression of *OsSHR2, OsRBOHC,* and *OsARM1* in the roots of primed rice plantlets (Fig 7). Among them, the *OsRBOHC* gene, which encodes for the respiratory burst oxidase homolog C, by regulating ROS production, contributes to the modification of cell walls, induction of programmed cell death, and ultimately, the formation of aerenchyma in rice roots allowing the rice plant to thrive in environments with limited oxygen availability. The finding that, aerenchyma contributes to the radial oxygen loss from roots to the rhizosphere and is particularly important in limiting the movement of As ions toward the root surface highlights the importance of *OsRBOHC* (Mei et al. 2012). Since this increased release of oxygen by aerenchyma into the rhizosphere could hinder As uptake into rice roots (Mei et al. 2012), the well-developed aerenchyma in the AvE-primed rice seedlings might explain the reduced uptake of As in the primed seedling by forming an oxidative shield around the growing roots and impeding the entry of As into the plants compared to unprimed ones under As stress. Taken together, these findings shed light on the probable intricate molecular responses elicited by AvE priming in the context of As stress, emphasizing the potential modulation of key genes involved in As transport and oxidative stress mitigation in rice seedlings.

At the agronomic level, the cumulative benefits of AvE priming translated into significant improvements in plant architecture, yield-related traits, and overall grain productivity under As stress. In our analysis, AvE priming promoted early flowering (7 to 10 days) or accelerated reproductive maturity. Since As stress affects rice most severely during the seedling and grain-filling stages, early flowering enables plants to complete grain setting before arsenic accumulates to reproductive toxicity levels within plant tissues. The observed variation in agronomic traits among rice plants highlights the importance of yield-contributing components in determining overall productivity. Flowering time is particularly critical because it influences the environmental conditions experienced during pollination and grain filling (Bheemanahalli et al. 2017). Plant height and seed size exhibited a moderate positive association with grain yield, suggesting that plants with optimal height may support improved panicle development and greater biomass accumulation (Singh et al. 2024). However, excessively tall plants may be more prone to lodging and do not necessarily result in higher yields. Tillering ability is one of the most important determinants of rice yield because each productive tiller bears a panicle. Previous studies have demonstrated that panicle number derived from primary tillers contributes significantly to overall grain yield in rice ( Huang et al. 2020). Panicle length and grain number per panicle also play crucial roles in yield determination. Longer panicles generally accommodate more spikelets, thereby potentially enhancing grain yield when grain-filling efficiency remains high. However, several studies have reported that grain number and grain-filling efficiency exert a stronger influence on yield than panicle length alone. The variation observed in 100-grain weight further indicates differences in assimilate partitioning and grain development (Singh et al. 2024). Overall, these findings demonstrate that agronomic traits are key determinants of rice productivity and can serve as important selection criteria in breeding and agronomic improvement programs (Singh et al. 2024; Shrestha et al. 2021) (Table 5).

Collectively, the findings of our study support a mechanistic model wherein AvE priming enhances arsenic tolerance reduces grain accumulation in rice through: (i) reinforcement of antioxidant defense systems, (ii) maintenance of osmotic and cellular homeostasis, (iii) enhanced metal chelation and vacuolar sequestration, (iv) modulation of stress-responsive gene expression, and (v) preservation of anatomical integrity (Fig. 9). This study represents one of the first reports demonstrating the efficacy of a weed-derived phytoextract as a seed priming agent for mitigating arsenic stress in rice. Unlike transgenic approaches, this strategy is simple, economical, environmentally sustainable, and readily adaptable for field conditions. The utilisation of *A. viridis*, an abundant and underutilised weed, further enhances its translational potential. In conclusion, AvE-mediated phytopriming offers a promising, non-transgenic solution for mitigating arsenic stress and reducing its accumulation in rice. Future investigations focusing on metabolite-specific functional validation, large-scale field trials, and long-term soil–plant interactions will be essential to fully harness the potential of this approach for sustainable agriculture in arsenic-affected regions.

**Figure 9.**
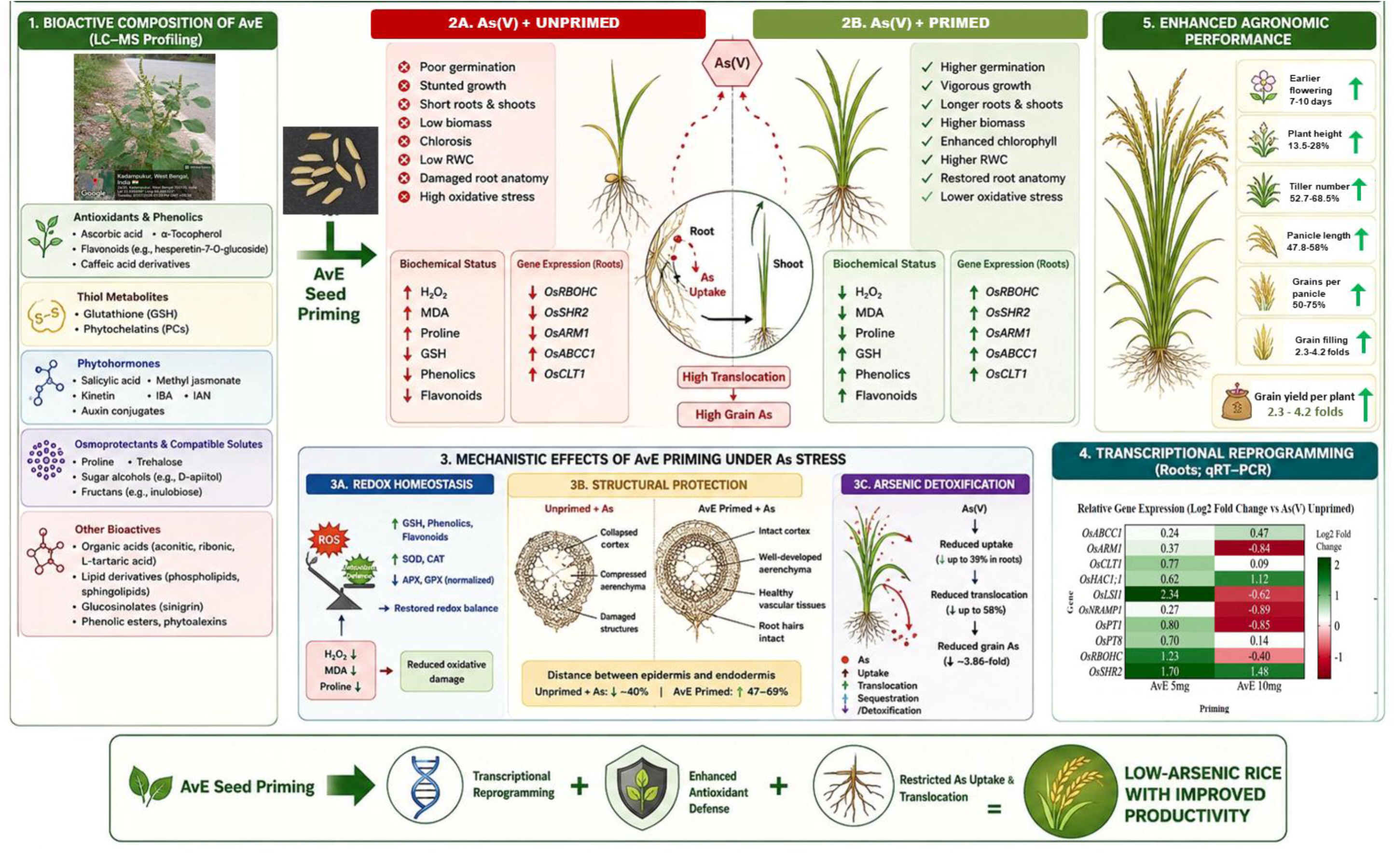
A schematic diagram depicting the overview of the possible pathway of AvE-mediated seed priming restricting As accumulation simultaneously improving As stress resilience in rice plants (the figure was generated using the AI tool ChatGPT using our research data as prompt).

## Acknowledgment

This work was supported by the Anusandhan National Research Foundation, Govt. of India, [grant CRG/2022/007083] and Department of Science and Technology and Biotechnology, Govt of West Bengal [grant 2296 (Sanc.)/STBT-13015/8/2024-WBSCST SEC] to SB. SP and SR acknowledge the Department of Science and Technology and Biotechnology, Govt of West Bengal and Council of Scientific and Industrial Research, respectively for providing fellowships. Amity University Kolkata is acknowledged for providing the infrastructural support. The authors are sincerely thankful to Dr. Maumita Bandopadhyay, Department of Botany, University of Calcutta for LC-MS analysis. The Authors sincerely acknowledge the Patent Information Centre, West Bengal State Council of Science and Technology, Department of Science and Technology and Biotechnology, Government of West Bengal for their support during intellectual property rights application (Indian Patent application No. 202631073698).

## Author contributions

**SP:** Methodology, Investigation, Data curation, Writing-Original draft preparation **SR:** Methodology, Data curation, Writing-Original draft preparation, **AB:** Investigation, Data curation, Writing-Original draft preparation **IDS:** Investigation, Data curation, Writing-Original draft preparation, **SC:** Investigation, Data curation **RS:** Supervision **ND:** Supervision, Writing-Reviewing and Editing **SB:** Conceptualization, Methodology, Supervision, Writing-Reviewing and Editing, Project administration, Funding acquisition

## Supplementary Material

**Supplementary Material 1:** Qualitative phytochemical characterisation of *Amaranthus viridis*.

**Supplementary Material 2:** Metabolic analysis of *Amaranthus viridis* by LC-MS profiling.

**Supplementary Material 3:** Effect of *Amaranthus viridis* extract (AvE) seed priming on germination, growth, physiological attributes, antioxidant defense, oxidative stress, and arsenic accumulation in rice seedlings exposed to As(III) stress. The figure illustrates the effects of seed priming with aqueous *Amaranthus viridis* extract (AvE 5 mg mL□¹ and AvE 10 mg mL□¹) on the morphophysiological, biochemical, and antioxidant responses of rice seedlings grown under sodium arsenite [As(III)] stress. Comparisons were made among (A) As(III) + unprimed, (B) As(III) + AvE 5 mg mL□¹, and (C) As(III) + AvE 10 mg mL□¹ treatments.

**Supplementary Material 4:** AvE seed priming alleviates As-induced anatomical alterations in the roots of 14-day-old (14 DAS) rice seedlings. Transverse sections of the roots showing (a) bright-field images (upper panel) and (b) EDAX images (lower panel) of Control, As(V) + unprimed, As(V) + AvE 5 mg mL□¹, and As(V) + AvE 10 mg mL□¹ treatments, illustrating the anatomical alterations induced by As stress and their alleviation following AvE seed priming.

**Supplementary Material 5:** List of oligonucleotide primers used for quantitative real-time PCR (qRT-PCR) analysis of stress-responsive and root morphological marker genes.

**Supplementary Material 6:** Relative expression analysis of stress-responsive and root morphological marker genes in rice roots by quantitative real-time PCR (qRT-PCR) under As(V) + unprimed, As(V) + AvE 5 mg mL□¹, and As(V) + AvE 10 mg mL□¹ treatments.

## Reference

Abbas, G., Murtaza, B., Bibi, I., Shahid, M., Niazi, N., Khan, M., Amjad, M., Hussain, M., & Natasha. (2018). Arsenic uptake, toxicity, detoxification, and speciation in plants: Physiological, biochemical, and molecular aspects. International Journal of Environmental Research and Public Health, 15(1), 59. 10.3390/ijerph15010059

Abdul Baki, A. A., & Anderson, J. D. (1973). Vigor determination in soybean seed by multiple criteria^1^. Crop Science, 13(6), 630–633. 10.2135/cropsci1973.0011183X001300060013x

Acharya, P., Jayaprakasha, G. K., Crosby, K. M., Jifon, J. L., & Patil, B. S. (2020). Nanoparticle-mediated seed priming improves germination, growth, yield, and quality of watermelons (Citrullus lanatus) at multi-locations in texas. Scientific Reports, 10(1), 5037. 10.1038/s41598-020-61696-7

Agati, G., Azzarello, E., Pollastri, S., & Tattini, M. (2012). Flavonoids as antioxidants in plants: Location and functional significance. Plant Science: An International Journal of Experimental Plant Biology, 196, 67–76. 10.1016/j.plantsci.2012.07.014

Ahuja, I., Kissen, R., & Bones, A. M. (2012). Phytoalexins in defense against pathogens. Trends in Plant Science, 17(2), 73–90. 10.1016/j.tplants.2011.11.002

Akpinar, A., & Cansev, A. (2024). Choline supplementation reduces cadmium uptake and alleviates cadmium toxicity in Solanum lycopersicum seedlings. BMC Plant Biology, 24(1), 977. 10.1186/s12870-024-05653-w

Ali, S., Abbas, Z., Seleiman, M. F., Rizwan, M., Yavaş, İ., Alhammad, B. A., Shami, A., Hasanuzzaman, M., & Kalderis, D. (2020). Glycine betaine accumulation, significance and interests for heavy metal tolerance in plants. Plants, 9(7), 896. 10.3390/plants9070896

Ali, U., Li, H., Wang, X., & Guo, L. (2018). Emerging roles of sphingolipid signaling in plant response to biotic and abiotic stresses. Molecular Plant, 11(11), 1328–1343. 10.1016/j.molp.2018.10.001

Analysis of Major, Minor and Trace Elements in Animal Tissue Samples with ICP-OES and ICP-MS, (October 2005), Standard Operation Procedure, Soil & Plant Analysis Laboratory, University of Wisconsin – Madison, https://uwlab.soils.wisc.edu/wp-content/uploads/sites/17/2015/08/animal_icp.pdf

Anderson, M. E. (1985). Determination of glutathione and glutathione disulfide in biological samples. Methods in Enzymology, 113, 548–555. 10.1016/s0076-6879(85)13073-9

Anjum, S. A., Tanveer, M., Hussain, S., Shahzad, B., Ashraf, U., Fahad, S., Hassan, W., Jan, S., Khan, I., Saleem, M. F., Bajwa, A. A., Wang, L., Mahmood, A., Samad, R. A., & Tung, S. A. (2016). Osmoregulation and antioxidant production in maize under combined cadmium and arsenic stress. Environmental Science and Pollution Research, 23(12), 11864–11875. 10.1007/s11356-016-6382-1

Arthur, K. L., Turner, M. A., Reynolds, J. C., & Creaser, C. S. (2017). Increasing peak capacity in nontargeted omics applications by combining full scan field asymmetric waveform ion mobility spectrometry with liquid chromatography–mass spectrometry. Analytical Chemistry, 89(6), 3452–3459. 10.1021/acs.analchem.6b04315

Awasthi, S., Chauhan, R., Srivastava, S., & Tripathi, R. D. (2017). The journey of arsenic from soil to grain in rice. Frontiers in Plant Science, 8, 1007. 10.3389/fpls.2017.01007

Bakry, B. A., Taha, M. H., Abdelgawad, Z. A., & Abdallah, M. M. S. (2014). The role of of three flax cultivars grown under saline soil conditions. Agricultural Sciences, 05(14), 1566–1575. 10.4236/as.2014.514168

Banerjee, S., Islam, J., Mondal, S., Saha, A., Saha, B., & Sen, A. (2023). Proactive attenuation of arsenic-stress by nano-priming: Zinc Oxide Nanoparticles in Vigna mungo (L.) Hepper trigger antioxidant defense response and reduce root-shoot arsenic translocation. Journal of Hazardous Materials, 446, 130735. 10.1016/j.jhazmat.2023.130735

Barnes, J. D., Balaguer, L., Manrique, E., Elvira, S., & Davison, A. W. (1992). A reappraisal of the use of DMSO for the extraction and determination of chlorophylls a and b in lichens and higher plants. Environmental and Experimental Botany, 32(2), 85–100. 10.1016/0098-8472(92)90034-Y

Barrs, H. D., & Weatherley, P. E. (1962). A re-examination of the relative turgidity technique for estimating water deficits in leaves. Australian Journal of Biological Sciences, 15(3), 413–428. 10.1071/bi9620413

Basra, S. M. A., Farooq, M., Tabassam, R., & Ahmad, N. (2005). Physiological and biochemical aspects of pre-sowing seed treatments in fine rice (Oryza sativa L.). Seed Science and Technology, 33(3), 623–628. 10.15258/sst.2005.33.3.09

Bates, L. S., Waldren, R. P., & Teare, I. D. (1973). Rapid determination of free proline for water-stress studies. Plant and Soil, 39(1), 205–207. 10.1007/BF00018060

Benkeblia, N. (2022). Insights on fructans and resistance of plants to drought stress. Frontiers in Sustainable Food Systems, 6, 827758. 10.3389/fsufs.2022.827758

Bheemanahalli, R., Sathishraj, R., Manoharan, M., Sumanth, H. N., Muthurajan, R., Ishimaru, T., & Krishna, J. S. V. (2017). Is early morning flowering an effective trait to minimize heat stress damage during flowering in rice? Field Crops Research, 203, 238–242. 10.1016/j.fcr.2016.11.011

Birla, D. S., Malik, K., Sainger, M., Chaudhary, D., Jaiwal, R., & Jaiwal, P. K. (2017). Progress and challenges in improving the nutritional quality of rice (oryza sativa l.). Critical Reviews in Food Science and Nutrition, 57(11), 2455–2481. 10.1080/10408398.2015.1084992

Brain, K. R., & Turner, T. D. (1975). The practical evaluation of phytopharmaceuticals. Bristol: Wright Sccintecnica, 81–82.

Castro, J. C., Castro, C. G., & Cobos, M. (2023). Genetic and biochemical strategies for regulation of L-ascorbic acid biosynthesis in plants through the L-galactose pathway. Frontiers in Plant Science, 14. 10.3389/fpls.2023.1099829

Chen, C. C., & Sung, J. M. (2001). Priming bitter gourd seeds with selenium solution enhances germinability and antioxidative responses under sub optimal temperature. Physiologia Plantarum, 111(1), 9–16. 10.1034/j.1399-3054.2001.1110102.x

Chobot, V., & Hadacek, F. (2009). Milieu-dependent pro- and antioxidant activity of juglone may explain linear and nonlinear effects on seedling development. Journal of Chemical Ecology, 35(3), 383–390. 10.1007/s10886-009-9609-5

Chowardhara, B., Borgohain, P., Saha, B., Awasthi, J. P., Moulick, D., & Panda, S. K. (2019). Phytotoxicity of Cd and Zn on three popular Indian mustard varieties during germination and early seedling growth. Biocatalysis and Agricultural Biotechnology, 21, 101349. 10.1016/j.bcab.2019.101349

Chowardhara, B., Saha, B., Borgohain, P., Awasthi, J. P., Kityania, S., & Panda, S. K. (2022). Effect of ethanol, putresciene and acetic acid on cadmium accumulation and toxicity in Indian mustard. South African Journal of Botany, 147, 42–52. 10.1016/j.sajb.2021.12.019

Clouse S.D. (2011). Brassinosteroid signal transduction: from receptor kinase activation to transcriptional networks regulating plant development and stress responses. The Plant Cell, 23(4), 1219–1230. 10.1105/tpc.111.084475

Cortleven, A., & Schmülling, T. (2015). Regulation of chloroplast development and function by cytokinin. Journal of Experimental Botany, 66(16), 4999–5013. 10.1093/jxb/erv132

Cortleven, A., Leuendorf, J. E., Frank, M., Pezzetta, D., Bolt, S., & Schmülling, T. (2019). Cytokinin action in response to abiotic and biotic stresses in plants. Plant, Cell & Environment, 42(3), 998–1018. 10.1111/pce.13494

Damodaran, S., & Strader, L. C. (2019). Indole 3-butyric acid metabolism and transport in arabidopsis thaliana. Frontiers in Plant Science, 10. 10.3389/fpls.2019.00851

Dao, T. T. H., Linthorst, H. J. M., & Verpoorte, R. (2011). Chalcone synthase and its functions in plant resistance. Phytochemistry Reviews, 10(3), 397–412. 10.1007/s11101-011-9211-7

Das, A., Majumder, S., Barman, S., Chatterjee, D., Mukhopadhyay, S., Ghosh, P., Pal, C. N., & Saha, G. (2021). Influence of basin-wide geomorphology on arsenic distribution in Nadia district. Environmental Research, 192, 110314. 10.1016/j.envres.2020.110314

Das, N., Bhattacharya, S., & Maiti, M. K. (2020). Biotechnological strategies to reduce arsenic content in rice. In S. Srivastava (Ed.), Arsenic in Drinking Water and Food (pp. 445–460). Springer Singapore. 10.1007/978-981-13-8587-2_18

Das, N., Bhattacharya, S., Bhattacharyya, S., & Maiti, M. K. (2017). Identification of alternatively spliced transcripts of rice phytochelatin synthase 2 gene OsPCS2 involved in mitigation of cadmium and arsenic stresses. Plant Molecular Biology, 94(1–2), 167–183. 10.1007/s11103-017-0600-1

Dewhirst, R. A., Lei, J., Afseth, C. A., Castanha, C., Wistrom, C. M., Mortimer, J. C., & Jardine, K. J. (2021). Are methanol-derived foliar methyl acetate emissions a tracer of acetate-mediated drought survival in plants? Plants, 10(2), 411. 10.3390/plants10020411

Ejiri, M., Fukao, T., Miyashita, T., & Shiono, K. (2021). A barrier to radial oxygen loss helps the root system cope with waterlogging-induced hypoxia. Breeding Science, 71(1), 40–50. 10.1270/jsbbs.20110

Eluwa, M. C. (1977). Studies on Gasteroclisus rhomboidalis (Boheman.) (Coleoptera: Curculionidae)—a pest of the African ‘spinach.’ Journal of Natural History, 11(4), 417–424. 10.1080/00222937700770331

Emamverdian, A., Ding, Y., Mokhberdoran, F., & Xie, Y. (2015). Heavy metal stress and some mechanisms of plant defense response. The Scientific World Journal, 2015, 1–18. 10.1155/2015/756120

Ende, W. V. D. (2013). Multifunctional fructans and raffinose family oligosaccharides. Frontiers in Plant Science, 4. 10.3389/fpls.2013.00247

Trease, E.C. and Evans, W.C. (2009) Pharmacognosy. 16th Edition, W.B. Saunders, Philadelphia, 365–650.

Felemban, A., Braguy, J., Zurbriggen, M. D., & Al-Babili, S. (2019). Apocarotenoids involved in plant development and stress response. Frontiers in Plant Science, 10. 10.3389/fpls.2019.01168

Feng, R., Wei, C., & Tu, S. (2013). The roles of selenium in protecting plants against abiotic stresses. Environmental and Experimental Botany, 87, 58–68. 10.1016/j.envexpbot.2012.09.002

Fernandez, O., Béthencourt, L., Quero, A., Sangwan, R. S., & Clément, C. (2010). Trehalose and plant stress responses: Friend or foe? Trends in Plant Science, 15(7), 409–417. 10.1016/j.tplants.2010.04.004

Fernie, A. R., Carrari, F., & Sweetlove, L. J. (2004). Respiratory metabolism: Glycolysis, the TCA cycle and mitochondrial electron transport. Current Opinion in Plant Biology, 7(3), 254–261. 10.1016/j.pbi.2004.03.007

Fitzpatrick, T. B. (2024). B vitamins: An update on their importance for plant homeostasis. Annual Review of Plant Biology, 75(1), 67–93. 10.1146/annurev-arplant-060223-025336

Flügel, F., Timm, S., Arrivault, S., Florian, A., Stitt, M., Fernie, A. R., & Bauwe, H. (2017). The photorespiratory metabolite 2-phosphoglycolate regulates photosynthesis and starch accumulation in arabidopsis. The Plant Cell, 29(10), 2537–2551. 10.1105/tpc.17.00256

Gao, M. (2020). Coronatine is more potent than jasmonates in regulating arabidopsis circadian clock. ASPB PLANT BIOLOGY 2020. ASPB PLANT BIOLOGY 2020. 10.46678/PB.20.1046369

Geng X., Cheng J., Gangadharan A., Mackey D. (2012). The coronatine toxin of Pseudomonas syringae is a multifunctional suppressor of Arabidopsis defense. Plant Cell, 24(11), 4763–4774. 10.1105/tpc.112.101402

Ghosh, S., Banerjee, S., & Sil, P. C. (2015). The beneficial role of curcumin on inflammation, diabetes and neurodegenerative disease: A recent update. Food and Chemical Toxicology, 83, 111–124. 10.1016/j.fct.2015.05.022

Gómez-Espinoza, O., Rojas-Villalta, D., Zúñiga-Pereira, A. M., Chacón-Díaz, C., Bravo, L. A., & Reyes-Díaz, M. (2025). Plant oxalate oxidases: Key enzymes in redox and stress regulation. Journal of Experimental Botany, 76(17), 4896–4909. 10.1093/jxb/eraf317

Grace, S. C. (2005). Phenolics as antioxidants. In N. Smirnoff (Ed.), Antioxidants and Reactive Oxygen Species in Plants (1st edn, pp. 141–168). Wiley. 10.1002/9780470988565.ch6

Han, X., & Yang, Y. (2021). Phospholipids in salt stress response. Plants, 10(10), 2204. 10.3390/plants10102204

Harborne, J. B. (1984). Phytochemical methods. Springer Netherlands. 10.1007/978-94-009-5570-7

Harris, D., Rashid, A., Miraj, G., Arif, M., & Yunas, M. (2008). ‘On-farm’ seed priming with zinc in chickpea and wheat in Pakistan. Plant and Soil, 306(1–2), 3–10. 10.1007/s11104-007-9465-4

Hasanuzzaman, M., Bhuyan, M. H. M. B., Raza, A., Hawrylak-Nowak, B., Matraszek-Gawron, R., Mahmud, J. A., Nahar, K., & Fujita, M. (2020). Selenium in plants: Boon or bane? Environmental and Experimental Botany, 178, 104170. 10.1016/j.envexpbot.2020.104170

Hayat, Q., Hayat, S., Irfan, Mohd., & Ahmad, A. (2010). Effect of exogenous salicylic acid under changing environment: A review. Environmental and Experimental Botany, 68(1), 14–25. 10.1016/j.envexpbot.2009.08.005

Heath, R. L., & Packer, L. (1968). Photoperoxidation in isolated chloroplasts. Archives of Biochemistry and Biophysics, 125(1), 189–198. 10.1016/0003-9861(68)90654-1

Hildebrandt, T. M. (2018). Synthesis versus degradation: Directions of amino acid metabolism during Arabidopsis abiotic stress response. Plant Molecular Biology, 98(1), 121–135. 10.1007/s11103-018-0767-0

Hiltunen, H.-M., Illarionov, B., Hedtke, B., Fischer, M., & Grimm, B. (2012). Arabidopsis RIBA proteins: Two out of three isoforms have lost their bifunctional activity in riboflavin biosynthesis. International Journal of Molecular Sciences, 13(11), 14086– 14105. 10.3390/ijms131114086

Hu, Y., Li, J., Lou, B., Wu, R., Wang, G., Lu, C., Wang, H., Pi, J., & Xu, Y. (2020). The role of reactive oxygen species in arsenic toxicity. Biomolecules, 10(2), 240. 10.3390/biom10020240

Huang, M., Shan, S., Cao, J., Fang, S., Tian, A., Liu, Y., Cao, F., Yin, X., & Zou, Y. (2020). Primary-tiller panicle number is critical to achieving high grain yields in machine-transplanted hybrid rice. Scientific Reports, 10(1), 2811. 10.1038/s41598-020-59751-4

Jiadkong, K., Fauzia, A. N., Yamaguchi, N., & Ueda, A. (2024). Exogenous riboflavin (Vitamin b2) application enhances salinity tolerance through the activation of its biosynthesis in rice seedlings under salinity stress. Plant Science, 339, 111929. 10.1016/j.plantsci.2023.111929

Khan, I., Awan, S. A., Rizwan, M., Ali, S., Zhang, X., & Huang, L. (2021). Arsenic behavior in soil-plant system and its detoxification mechanisms in plants: A review. Environmental Pollution, 286, 117389. 10.1016/j.envpol.2021.117389

Kim, J.-M., To, T. K., Matsui, A., Tanoi, K., Kobayashi, N. I., Matsuda, F., Habu, Y., Ogawa, D., Sakamoto, T., Matsunaga, S., Bashir, K., Rasheed, S., Ando, M., Takeda, H., Kawaura, K., Kusano, M., Fukushima, A., Endo, T. A., Kuromori, T., … Seki, M. (2017). Acetate-mediated novel survival strategy against drought in plants. Nature Plants, 3(7), 17097. 10.1038/nplants.2017.97

Korasick, D. A., Enders, T. A., & Strader, L. C. (2013). Auxin biosynthesis and storage forms. Journal of Experimental Botany, 64(9), 2541–2555. 10.1093/jxb/ert080

Lei, S., Rossi, S., Yang, Z., Yu, J., & Huang, B. (2024). Metabolic regulation of 5-oxoproline for enhanced heat tolerance in perennial ryegrass. Stress Biology, 4(1), 46. 10.1007/s44154-024-00175-9

Linster, C. L., & Clarke, S. G. (2008). L-Ascorbate biosynthesis in higher plants: The role of VTC2. Trends in Plant Science, 13(11), 567–573. 10.1016/j.tplants.2008.08.005

López□Lara, I. M., Nogales, J., Pech Canul, Á., Calatrava Morales, N., Bernabéu Roda, L. M., Durán, P., Cuéllar, V., Olivares, J., Alvarez, L., Palenzuela Bretones, D., Romero, M., Heeb, S., Cámara, M., Geiger, O., & Soto, M. J. (2018). 2 Tridecanone impacts surface associated bacterial behaviours and hinders plant–bacteria interactions. Environmental Microbiology, 20(6), 2049–2065. 10.1111/1462-2920.14083

Lunn, J. E., Delorge, I., Figueroa, C. M., Van Dijck, P., & Stitt, M. (2014). Trehalose metabolism in plants. The Plant Journal, 79(4), 544–567. 10.1111/tpj.12509

Lv, S., Tai, F., Guo, J., Jiang, P., Lin, K., Wang, D., Zhang, X., & Li, Y. (2020). Phosphatidylserine synthase from salicornia europaea is involved in plant salt tolerance by regulating plasma membrane stability. Plant and Cell Physiology, 62(1), 66–79. 10.1093/pcp/pcaa141

Ma, X., Sharifan, H., Dou, F., & Sun, W. (2020). Simultaneous reduction of arsenic (As) and cadmium (Cd) accumulation in rice by zinc oxide nanoparticles. Chemical Engineering Journal, 384, 123802. 10.1016/j.cej.2019.123802

Maeda, H., & Dudareva, N. (2012). The shikimate pathway and aromatic amino acid biosynthesis in plants. Annual Review of Plant Biology, 63(1), 73–105. 10.1146/annurev-arplant-042811-105439

Mahajan, G., Kumar, V., & Chauhan, B. S. (2017). Rice production in india. In B. S. Chauhan, K. Jabran, & G. Mahajan (Eds.), Rice Production Worldwide (pp. 53–91). Springer International Publishing. 10.1007/978-3-319-47516-5_3

Mathur, P., Tripathi, D. K., Baluška, F., & Mukherjee, S. (2022). Auxin-mediated molecular mechanisms of heavy metal and metalloid stress regulation in plants. Environmental and Experimental Botany, 196, 104796. 10.1016/j.envexpbot.2022.104796

Meents, A. K., Chen, S.-P., Reichelt, M., Lu, H.-H., Bartram, S., Yeh, K.-W., & Mithöfer, A. (2019). Volatile DMNT systemically induces jasmonate-independent direct anti-herbivore defense in leaves of sweet potato (Ipomoea batatas) plants. Scientific Reports, 9(1), 17431. 10.1038/s41598-019-53946-0

Mei, X. Q., Wong, M. H., Yang, Y., Dong, H. Y., Qiu, R. L., & Ye, Z. H. (2012). The effects of radial oxygen loss on arsenic tolerance and uptake in rice and on its rhizosphere. Environmental Pollution, 165, 109–117. 10.1016/j.envpol.2012.02.018

Moulick, D., Ghosh, D., & Chandra Santra, S. (2016). Evaluation of effectiveness of seed priming with selenium in rice during germination under arsenic stress. Plant Physiology and Biochemistry, 109, 571–578. 10.1016/j.plaphy.2016.11.004

Mridha, D., Paul, I., De, A., Ray, I., Das, A., Joardar, M., Chowdhury, N. R., Bhadoria, P. B. S., & Roychowdhury, T. (2021). Rice seed (Ir64) priming with potassium humate for improvement of seed germination, seedling growth and antioxidant defense system under arsenic stress. Ecotoxicology and Environmental Safety, 219, 112313. 10.1016/j.ecoenv.2021.112313

Munné-Bosch, S. (2005). The role of α -tocopherol in plant stress tolerance. Journal of Plant Physiology, 162(7), 743–748. 10.1016/j.jplph.2005.04.022

Murugaiyan, V., Zeibig, F., Anumalla, M., Siddiq, S. A., Frei, M., Murugaiyan, J., & Ali, J. (2021). Arsenic stress responses and accumulation in rice. In J. Ali & S. H. Wani (Eds.), Rice Improvement (pp. 281–313). Springer International Publishing. 10.1007/978-3-030-66530-2_9

Nakamoto, M., Kunimura, K., Suzuki, J., & Kodera, Y. (2019). Antimicrobial properties of hydrophobic compounds in garlic: Allicin, vinyldithiin, ajoene and diallyl polysulfides (Review). Experimental and Therapeutic Medicine. 10.3892/etm.2019.8388

Patade, V. Y., Bhargava, S., & Suprasanna, P. (2009). Halopriming imparts tolerance to salt and PEG induced drought stress in sugarcane. Agriculture, Ecosystems & Environment, 134(1–2), 24–28. 10.1016/j.agee.2009.07.003

Rahman, M. A., Rahman, M. M., & Naidu, R. (2014). Arsenic in rice. In Wheat and Rice in Disease Prevention and Health (pp. 365–375). Elsevier. 10.1016/B978-0-12-401716-0.00028-3

Ray, I., Mridha, D., Sarkar, J., Joardar, M., Das, A., Chowdhury, N. R., De, A., Acharya, K., & Roychowdhury, T. (2022). Application of potassium humate to reduce arsenic bioavailability and toxicity in rice plants (Oryza sativa L.) during its course of germination and seedling growth. Environmental Pollution, 313, 120066. 10.1016/j.envpol.2022.120066

Reyad-ul-Ferdous, Md. (2015). Present biological status of potential medicinal plant of amaranthus viridis: A comprehensive review. American Journal of Clinical and Experimental Medicine, 3(5), 12. 10.11648/j.ajcem.s.2015030501.13

Sarkar, A., & Paul, B. (2016). The global menace of arsenic and its conventional remediation—A critical review. Chemosphere, 158, 37–49. 10.1016/j.chemosphere.2016.05.043

Schat, H., Sharma, S. S., & Vooijs, R. (1997). Heavy metal□induced accumulation of free proline in a metal□tolerant and a nontolerant ecotype of Silene vulgaris. Physiologia Plantarum, 101(3), 477–482. 10.1111/j.1399-3054.1997.tb01026.x

Schmo ger, M. E. V., Oven, M., & Grill, E. (2000). Detoxification of arsenic by phytochelatins in plants. Plant Physiology, 122(3), 793–802. 10.1104/pp.122.3.793

Shabbir, A., Shah, A. A., Usman, S., Ahmed, S., Kaleem, M., Shafique, S., & Gatasheh, M. K. (2025). Efficacy of malic and tartaric acid in mitigation of cadmium stress in Spinacia oleracea L. via modulations in physiological and biochemical attributes. Scientific Reports, 15(1), 3366. 10.1038/s41598-025-85896-1

Shaji, E., Santosh, M., Sarath, K. V., Prakash, P., Deepchand, V., & Divya, B. V. (2021). Arsenic contamination of groundwater: A global synopsis with focus on the Indian Peninsula. Geoscience Frontiers, 12(3), 101079. 10.1016/j.gsf.2020.08.015

Sharma, A., Shahzad, B., Rehman, A., Bhardwaj, R., Landi, M., & Zheng, B. (2019). Response of phenylpropanoid pathway and the role of polyphenols in plants under abiotic stress. Molecules, 24(13), 2452. 10.3390/molecules24132452

Shrestha, J., Subedi, S., Singh Kushwaha, U. K., & Maharjan, B. (2021). Evaluation of growth and yield traits in rice genotypes using multivariate analysis. Heliyon, 7(9), e07940. 10.1016/j.heliyon.2021.e07940

Shri, M., Kumar, S., Chakrabarty, D., Trivedi, P. K., Mallick, S., Misra, P., Shukla, D., Mishra, S., Srivastava, S., Tripathi, R. D., & Tuli, R. (2009). Effect of arsenic on growth, oxidative stress, and antioxidant system in rice seedlings. Ecotoxicology and Environmental Safety, 72(4), 1102–1110. 10.1016/j.ecoenv.2008.09.022

Singh, P., Kumar, A., Singh, T., Anto, S., Indoliya, Y., Tiwari, P., Behera, S. K., & Chakrabarty, D. (2024). Targeting OsNIP3;1 via CRISPR/Cas9: A strategy for minimizing arsenic accumulation and boosting rice resilience. Journal of Hazardous Materials, 471, 134325. 10.1016/j.jhazmat.2024.134325

Smirnoff, N. (2011). Vitamin c. In Advances in Botanical Research (Vol. 59, pp. 107–177). Elsevier. 10.1016/B978-0-12-385853-5.00003-9

Smits, J. E., Krohn, R. M., Akhtar, E., Hore, S. K., Yunus, Md., Vandenberg, A., & Raqib, R. (2019). Food as medicine: Selenium enriched lentils offer relief against chronic arsenic poisoning in Bangladesh. Environmental Research, 176, 108561. 10.1016/j.envres.2019.108561

So, M. J., & Cho, E. J. (2014). Phloroglucinol attenuates free radical-induced oxidative stress. Preventive Nutrition and Food Science, 19(3), 129–135. 10.3746/pnf.2014.19.3.129

Song, W.-Y., Yamaki, T., Yamaji, N., Ko, D., Jung, K.-H., Fujii-Kashino, M., An, G., Martinoia, E., Lee, Y., & Ma, J. F. (2014). A rice ABC transporter, OsABCC1, reduces arsenic accumulation in the grain. Proceedings of the National Academy of Sciences, 111(44), 15699–15704. 10.1073/pnas.1414968111

Srivastava, S., & Sharma, Y. K. (2014). Arsenic induced changes in growth and metabolism of black gram seedlings (Vigna mungo l.) and the role of phosphate as an ameliorating agent. Environmental Processes, 1(4), 431–445. 10.1007/s40710-014-0035-5

Srivastava, S., Sinha, P., & Sharma, Y. K. (2017). Status of photosynthetic pigments, lipid peroxidation and anti-oxidative enzymes in Vigna mungo in presence of arsenic. Journal of Plant Nutrition, 40(3), 298–306. 10.1080/01904167.2016.1240189

Szabados, L., & Savouré, A. (2010). Proline: A multifunctional amino acid. Trends in Plant Science, 15(2), 89–97. 10.1016/j.tplants.2009.11.009

Timm, S., Florian, A., Jahnke, K., Nunes-Nesi, A., Fernie, A. R., & Bauwe, H. (2011). The hydroxypyruvate-reducing system in Arabidopsis: Multiple enzymes for the same end. Plant Physiology, 155(2), 694–705. 10.1104/pp.110.166538

Vardhini, B. V., & Anjum, N. A. (2015). Brassinosteroids make plant life easier under abiotic stresses mainly by modulating major components of antioxidant defense system. Frontiers in Environmental Science, 2. 10.3389/fenvs.2014.00067

Varma, P. (2017). An overview of rice economy. In P. Varma, Rice Productivity and Food Security in India (pp. 7–28). Springer Singapore. 10.1007/978-981-10-3692-7_2

Velikova, V., Yordanov, I., & Edreva, A. (2000). Oxidative stress and some antioxidant systems in acid rain-treated bean plants: Protective role of exogenous polyamines. Plant Science, 151(1), 59–66. 10.1016/S0168-9452(99)00197-1

Vinoth, L., & Manivasagaperumal, R. (2012). Phytochemical analysis and antibacterial activity of moringa oleifera lam. https://www.semanticscholar.org/paper/PHYTOCHEMICAL-ANALYSIS-AND-ANTIBACTERIAL-ACTIVITY-Vinoth-Manivasagaperumal/044674d12e940166d5a6e56c3a0c113fcd0e00e3

Wasternack, C., & Song, S. (2017). Jasmonates: Biosynthesis, metabolism, and signaling by proteins activating and repressing transcription. Journal of Experimental Botany, 68(6), 1303–1321. 10.1093/jxb/erw443

Watanabe, S., Matsumoto, M., Hakomori, Y., Takagi, H., Shimada, H., & Sakamoto, A. (2014). The purine metabolite allantoin enhances abiotic stress tolerance through synergistic activation of abscisic acid metabolism. *Plant*, Cell & Environment, 37(4), 1022–1036. 10.1111/pce.12218

Williams, P. N., Villada, A., Deacon, C., Raab, A., Figuerola, J., Green, A. J., Feldmann, J., & Meharg, A. A. (2007). Greatly enhanced arsenic shoot assimilation in rice leads to elevated grain levels compared to wheat and barley. Environmental Science & Technology, 41(19), 6854–6859. 10.1021/es070627i

Yadav, P. V., Kumari, M., & Ahmed, Z. (2011). Chemical seed priming as a simple technique to impart cold and salt stress tolerance in capsicum. Journal of Crop Improvement, 25(5), 497–503. 10.1080/15427528.2011.587139

Yan, S., Wu, F., Zhou, S., Yang, J., Tang, X., & Ye, W. (2021). Zinc oxide nanoparticles alleviate the arsenic toxicity and decrease the accumulation of arsenic in rice (Oryza sativa L.). BMC Plant Biology, 21(1), 150. 10.1186/s12870-021-02929-3

Yang, J., Gao, M., Hu, H., Ding, X., Lin, H., Wang, L., Xu, J., Mao, C., Zhao, F., & Wu, Z. (2016). OsCLT1, a CRT like transporter 1, is required for glutathione homeostasis and arsenic tolerance in rice. New Phytologist, 211(2), 658–670. 10.1111/nph.13908

Yu, Y., Kou, M., Gao, Z., Liu, Y., Xuan, Y., Liu, Y., Tang, Z., Cao, Q., Li, Z., & Sun, J. (2019). Involvement of phosphatidylserine and triacylglycerol in the response of sweet potato leaves to salt stress. Frontiers in Plant Science, 10, 1086. 10.3389/fpls.2019.01086

Zarinkamar, F., Rezayian, M., & Medhat, R. (2022). Increase of trigonelline in trigonella persica plant under drought stress. Journal of Botanical Research, 4(2), 19–25. 10.30564/jbr.v4i2.4512

Zhao, F.-J., McGrath, S. P., & Meharg, A. A. (2010). Arsenic as a food chain contaminant: Mechanisms of plant uptake and metabolism and mitigation strategies. Annual Review of Plant Biology, 61(1), 535–559. 10.1146/annurev-arplant-042809-112152

Zhao, J., & Hu, J. (2023). Melatonin: Current status and future perspectives in horticultural plants. Frontiers in Plant Science, 14, 1140803. 10.3389/fpls.2023.1140803

Zhishen, J., Mengcheng, T., & Jianming, W. (1999). The determination of flavonoid contents in mulberry and their scavenging effects on superoxide radicals. Food Chemistry, 64(4), 555–559. 10.1016/S0308-8146(98)00102-2

